# Differences in spike generation instead of synaptic inputs determine the feature selectivity of two retinal cell types

**DOI:** 10.1101/2021.10.19.464988

**Authors:** Sophia Wienbar, Gregory William Schwartz

**Affiliations:** Department of Ophthalmology, Feinberg School of Medicine, Northwestern University, Chicago, IL, USA; Northwestern University Interdepartmental Neuroscience Program, Northwestern University, Evanston, IL, USA; Department of Physiology, Feinberg School of Medicine, Northwestern University, Chicago, IL, USA; Department of Neurobiology, Weinberg College of Arts and Sciences, Northwestern University, Evanston, IL, USA

## Abstract

The output of spiking neurons depends both on their synaptic inputs and on their intrinsic properties. Retinal ganglion cells (RGCs), the spiking projection neurons of the retina, comprise over 40 different types in mice and other mammals, each tuned to different features of visual scenes. The circuits providing synaptic input to different RGC types to drive feature selectivity have been studied extensively, but there has been substantially less research aimed at understanding how the intrinsic properties of RGCs differ and how those differences impact feature selectivity. Here, we introduce an RGC type in the mouse, the Bursty Suppressed-by-Contrast (bSbC) RGC, whose contrast selectivity is shaped by its intrinsic properties. Surprisingly, when we compare the bSbC RGC to the OFF sustained alpha (OFFsA) RGC that receives similar synaptic input, we find that the two RGC types exhibit starkly different responses to an identical stimulus. We identified spike generation as the key intrinsic property behind this functional difference; the bSbC RGC undergoes depolarization block in conditions where the OFFsA RGC maintains a high spike rate. Pharmacological experiments, imaging, and compartment modeling demonstrate that these differences in spike generation are the result of differences in voltage-gated sodium channel conductances. Our results demonstrate that differences in intrinsic properties allow these two RGC types to detect and relay distinct features of an identical visual stimulus to the brain.

## Introduction

The spikes of retinal ganglion cells (RGCs) convey all visual information to the brain. Our understanding of the typology of RGCs is most advanced in mice where they have been classified into more than 40 types based on differences in light responses, dendritic morphology, and gene expression (Bae et al., 2018; Goetz et al.; Tran et al., 2019). Each RGC type’s unique pattern of connectivity with upstream interneurons (bipolar and amacrine cells) plays a large role in determining the feature selectivity of its spike output, and many studies have recorded the synaptic inputs to RGCs in an effort to understand their light responses (Antinucci et al., 2016; Jacoby and Schwartz, 2017; Jacoby et al., 2015; Johnson et al., 2018; Kim et al., 2015; Vaney et al., 2012). Establishing differences in intrinsic properties is another mechanism by which the retina might expand the feature selectivity of the RGC population. However, little is known about the differences in the intrinsic properties of different RGC types and how these differences might contribute to feature selectivity.

Intrinsic properties can be conceptually divided into passive properties, like cellular morphology, membrane resistance, and capacitance; and active properties, that encompass properties of voltage-gated ion channels. Most links between the intrinsic properties of RGCs and their feature selectivity have focused on dendritic integration. Modeling and experimental studies have shown that the morphology of RGC dendritic arbors affects how they integrate synaptic input over visual space (Fohlmeister and Miller, 1997; Johnston and Lagnado, 2015; Koch et al., 1982; Ran et al., 2020; Stuart and Spruston, 2015), and active conductances in the dendrites of certain RGCs affect spatial integration (Abbas et al., 2013) and can lead to dendritic spikes that play a role in direction selectivity (Brombas et al., 2017; Oesch et al., 2005; Sivyer and Williams, 2013; Trenholm et al., 2014).

Following dendritic integration, spike generation at the axon initial segment (AIS) also depends on the different intrinsic properties of RGCs, leading to a wide diversity in the number, timing, and waveforms of action potentials among RGC types depolarized with the same somatic current injections (O’Brien et al., 2002; Wong et al., 2012). One profound way in which the properties of spike generation can influence information transmission in a neuron is by controlling susceptibility to depolarization block, a mechanism by which neurons cease firing during prolonged depolarization due to inactivation (block) of voltage-gated sodium channels. A modeling study demonstrated that the differential responses of rat RGCs to electrical stimulation arise from differential susceptibility to depolarization block and that active conductances and AIS length were the key variables controlling susceptibility (Kameneva et al., 2016). Recent work has shown that the biophysical properties controlling spike generation differ systematically among M1 intrinsically photosensitive (ip) RGCs, controlling their susceptibility to depolarization block to shape their light-sensitivity profiles (Emanuel et al., 2017; Milner and Do, 2017). It is not known whether depolarization block is used in other RGC types to shape feature selectivity and, more generally, whether the properties of spike generation in non-ipRGCs contribute in important ways to how they respond to visual stimuli.

We investigated the contribution of intrinsic properties to how two different RGC types encode a fundamental feature of a visual scene: contras, defined as the percent change in luminance from a mean light level. OFF sustained alpha (OFFsA) RGCs (homologous to OFF delta RGCs in rabbit and guinea pig (Krieger et al., 2017; Peichl, 1989; Rockhill et al., 2002; Roska et al., 2006)) are extremely sensitive to contrast, (Manookin et al., 2008). Their high baseline firing rate decreases for positive contrasts and increases for negative contrasts, and the high gain of the contrast response function depends on a push-pull motif of excitation and inhibition (Homann and Freed, 2017; Manookin et al., 2008). Here, we introduce the Bursty Suppressed-by-Contrast (bSbC) RGC and show that, while it has similar synaptic inputs to the OFFsA RGC, its spike response is qualitatively different; it decreases its baseline firing rate for both positive and negative contrasts, thus falling into the class of SbC RGC types (Jacoby and Schwartz, 2018; Jacoby et al., 2015; Tien et al., 2015). This cell decreases its firing rate to negative contrast as a result of depolarization block. Through a variety of methods, including electrophysiology, imaging, and compartment modeling, we demonstrate that the difference in how OFFsA and bSbC RGCs represent contrast is controlled not by different synaptic inputs but instead by differences in the properties of spike generation.

## Results

In a whole-mount *ex vivo* preparation of the mouse retina, we identified a novel RGC type and named it the Bursty Suppressed-by-Contrast (bSbC) RGC. Its key features included burst firing with irregular spike amplitudes and an SbC contrast response function: a decrease in baseline spike rate for both positive and negative contrast. Though it fell into the class of SbC RGCs functionally, the morphology and synaptic input to the bSbC RGC was much more similar to the well-known OFF sustained alpha (OFFsA) RGC (Murphy and Rieke, 2008, 2011; Pang et al., 2003), than it was to the two previously identified SbC RGCs in the mouse (Jacoby et al., 2015, 2018; Tien et al., 2015). Thus, throughout this work we compared bSbC RGCs to OFFsA RGCs.

### OFFsA and bSbC are distinct RGC types with different contrast response functions

We identified RGCs in whole-mounts of the wild-type mouse retina by their light responses and filled many of them with Neurobiotin or a fluorescent dye for morphological analysis (see **Methods** and (Goetz et al.)). The OFFsA RGC has been classified morphologically by its large soma and wide dendritic arbor in sublamina 1 of the inner plexiform layer (IPL), molecularly by its reactivity for the alpha RGC marker neurofilament protein SMI-32, and functionally by its high baseline firing rate and sustained OFF response (Bleckert et al., 2014; Murphy and Rieke, 2011; Pang et al., 2003). The bSbC RGC had several similarities to the OFFsA RGC, but also several differences that supported its classification as a distinct type (**Figure 1**). Morphologically, like the OFFsA, we found that the bSbC RGC was monostratified in the outer portion of the IPL and had a relatively large soma (**Figure 1A-C**). While these two RGC types had similar dendritic areas (**Figure 1G**, p = 0.35), bSbC RGCs had significantly smaller somas than OFFsA RGCs (**Figure 1F**, p = 0.010). The stratification patterns in the IPL had subtle differences that aligned with morphological types in the Eyewire museum (Bae et al., 2018). The OFFsA RGC matched type 1wt, confirming its functional assignment, and the bSbC RGC matched type 2o, which had no previous functional match (**Figure 1C**). The soma sizes of these two RGC types were also consistent with the electron microscopy data; type 2o has the fourth-largest soma after the three alpha RGCs (Bae et al., 2018). All bSbC RGC somas that we tested were negative for the alpha RGC marker SMI-32 (N = 4) while all OFFsA RGC somas were SMI-32 positive (N = 4) demonstrating a significant molecular difference (**Figure 1A**,**B**, p = 0.014, Fisher’s exact test). Finally, bSbC RGCs exhibited both a lower baseline firing rate (**Figure 1H**, p = 1.0 × 10^−07^) and more variability in spike amplitude than OFFsA RGCs (**Figure 1I**, p = 2.8 × 10^−14^).

**Figure 1.**
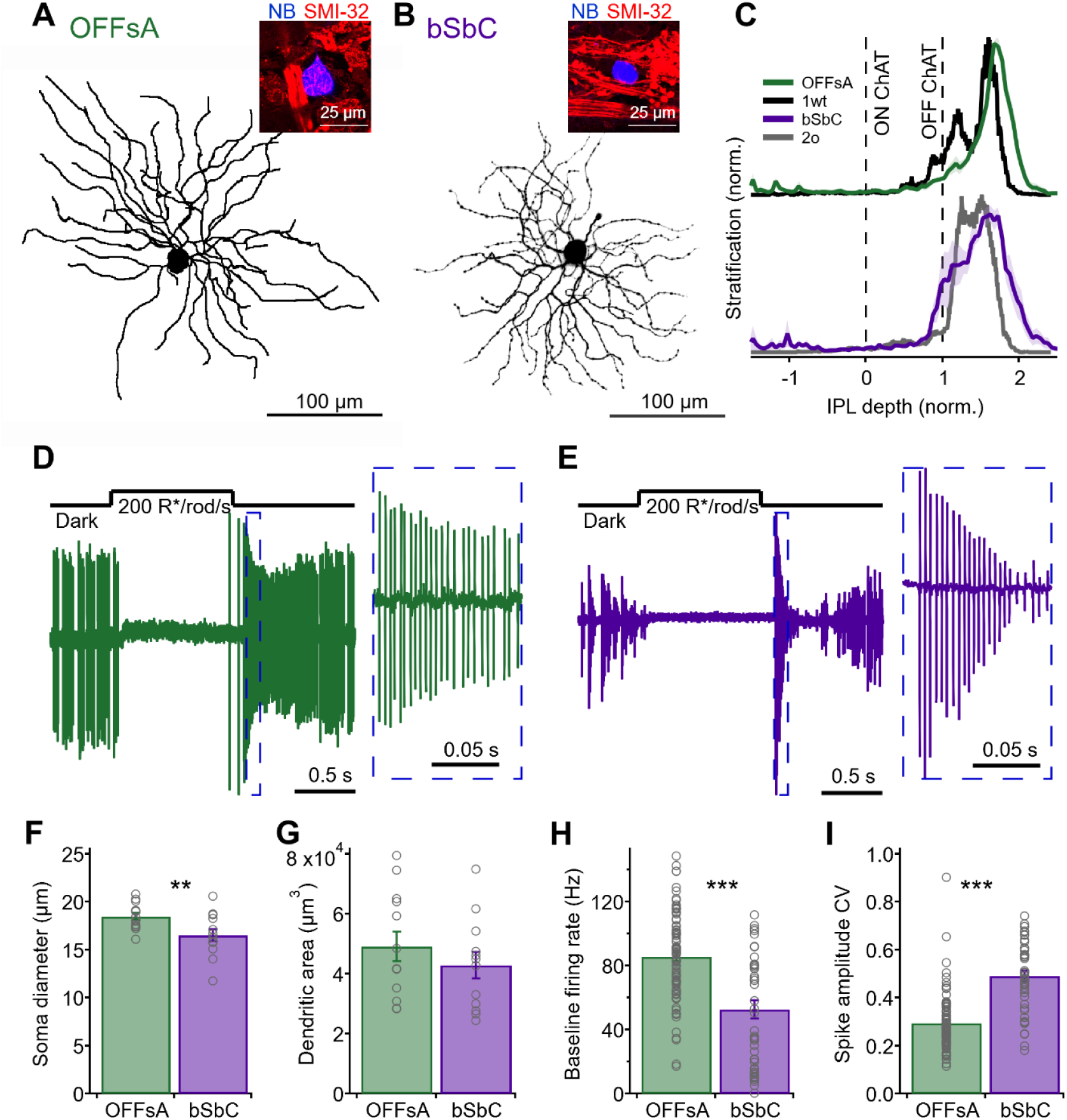
The OFFsA and bSbC are distinct RGC types. **(A, B)** *En face* view of an OFFsA **(A)** and a bSbC **(B)** dendritic arbor imaged using 2-photon. Insets show a Neurobiotin labeled soma of each RGC type (*blue*) and SMI-32 immunoreactivity (*red*). **(C)** Dendritic stratification for the OFFsA (*green*) as compared to the Eyewire type 1wt (*black*) and the bSbC (*purple*) as compared to the Eyewire type 2o (*gray*). Dotted lines refer to the ON and OFF ChAT (choline acetyltransferase) bands used to determine the stratification (see **Methods**). **(D, E)** Light step responses for an OFFsA **(D)** and a bSbC **(E)** measured in loose patch configuration. Insets show a magnified view of the OFF response. **(F)** Soma diameter of the OFFsA (N = 14) and bSbC (N = 13), p = 0.010. **(G)** Convex dendritic area of the OFFsA (N = 13) and bSbC (N = 12), p = 0.35. **(H)** Baseline firing rate in darkness of the OFFsA (N = 86) and bSbC (N = 47), p = 1.0 × 10^−07^. **(I)** Coefficient of variation (CV) of the baseline firing spike amplitudes in darkness of the OFFsA (N = 86) and bSbC (N = 47), p = 2.8 × 10^−14^. ** indicates a p-value < 0.01, *** indicates a p-value < 0.001.

Contrast sensitivity was different in OFFsA and bSbC RGCs, revealing a key functional distinction between these two cell types. In cell-attached recordings, OFFsA RGCs had a high baseline firing rate (86 ± 3 Hz, N = 86, **Figure 1H**), and they encoded small contrasts nearly linearly with high gain (**Figure 2A**). The baseline firing rate in bSbC RGCs (53 ± 6 Hz, N = 47, **Figure 1H**) decreased for both positive and negative contrasts (**Figure 2A**). Therefore, the OFFsA contrast response is classified as OFF polarity while the bSbC is classified as a suppressed-by-contrast cell. A closer examination of the spike responses of these RGCs to contrast steps revealed that the key functional difference occurs at negative contrasts where OFFsA RGCs fire at a high rate while bSbCs RGCs pause their maintained spiking (**Figure 2B**). Whole-cell current-clamp recordings provided an opportunity to measure subthreshold voltage changes in the two cell types during these light responses. These recordings confirmed the cell-attached contrast response measurements (**Figure 2C**), but they also provided additional insights into the source of this difference. As in OFFsA RGCs, positive contrasts elicited hyperpolarizations in bSbCs (**Figure 2D**). For negative contrasts, spike increases in OFFsA and spike decreases in bSbC RGCs were both accompanied by depolarizations (**Figure 2D**). While OFFsA RGCs maintained a high spike rate throughout the negative contrast step, bSbC RGCs went into depolarization block, preventing action potentials from being generated. Thus, OFFsA and bSbC RGCs have different contrast response functions that appear to be mediated by different responses to depolarization at negative contrast.

**Figure 2.**
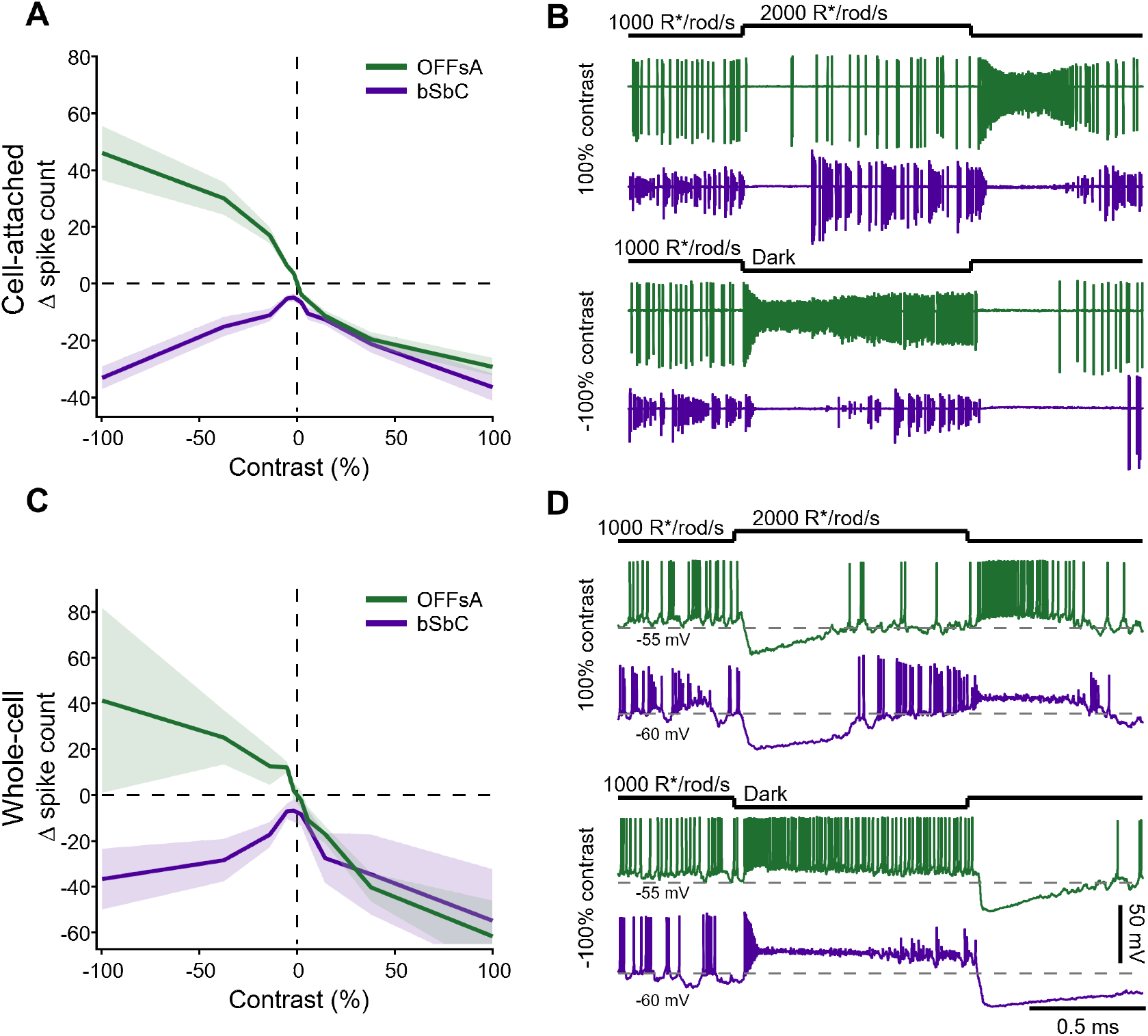
OFFsA and bSbC RGCs have different contrast response functions. **(A)** Contrast response functions in cell-attached recordings of OFFsA (*green*) and bSbC (*purple*) RGCs. Error bars are SEM across cells (N = 37 OFFsA, 15 bSbC). **(B)** Representative cell-attached traces of OFFsA (*green*) and bSbC (*purple*) RGCs responding to a +100% (top) and a -100% (bottom) contrast step. (N = 3 OFFsA, 4 bSbC) **(C)** Same as **(A)** but for whole-cell current clamp recordings. **(D)** Same as **(B)** but for whole-cell current clamp recordings.

### Synaptic inputs in OFFsA and bSbC RGCs are functionally interchangeable

To begin to disentangle the relative contributions of synaptic inputs and intrinsic properties on the contrast response functions, we recorded synaptic inputs to OFFsA and bSbC RGCs. We voltage clamped cells with a cesium based internal (see **Methods**) to isolate synaptic currents during the same contrast step stimuli we used to measure spikes (**Figure 3**). Both RGC types showed a push-pull pattern of increased excitation and decreased inhibition for negative contrast and the opposite changes for positive contrast (**Figure 3C**,**F**) as expected for the OFFsA (Homann and Freed, 2017; Manookin et al., 2008). The similarity in synaptic input was incongruous with the very different spike output in the two cell types for the same light stimuli (see **Figure 2**).

**Figure 3.**
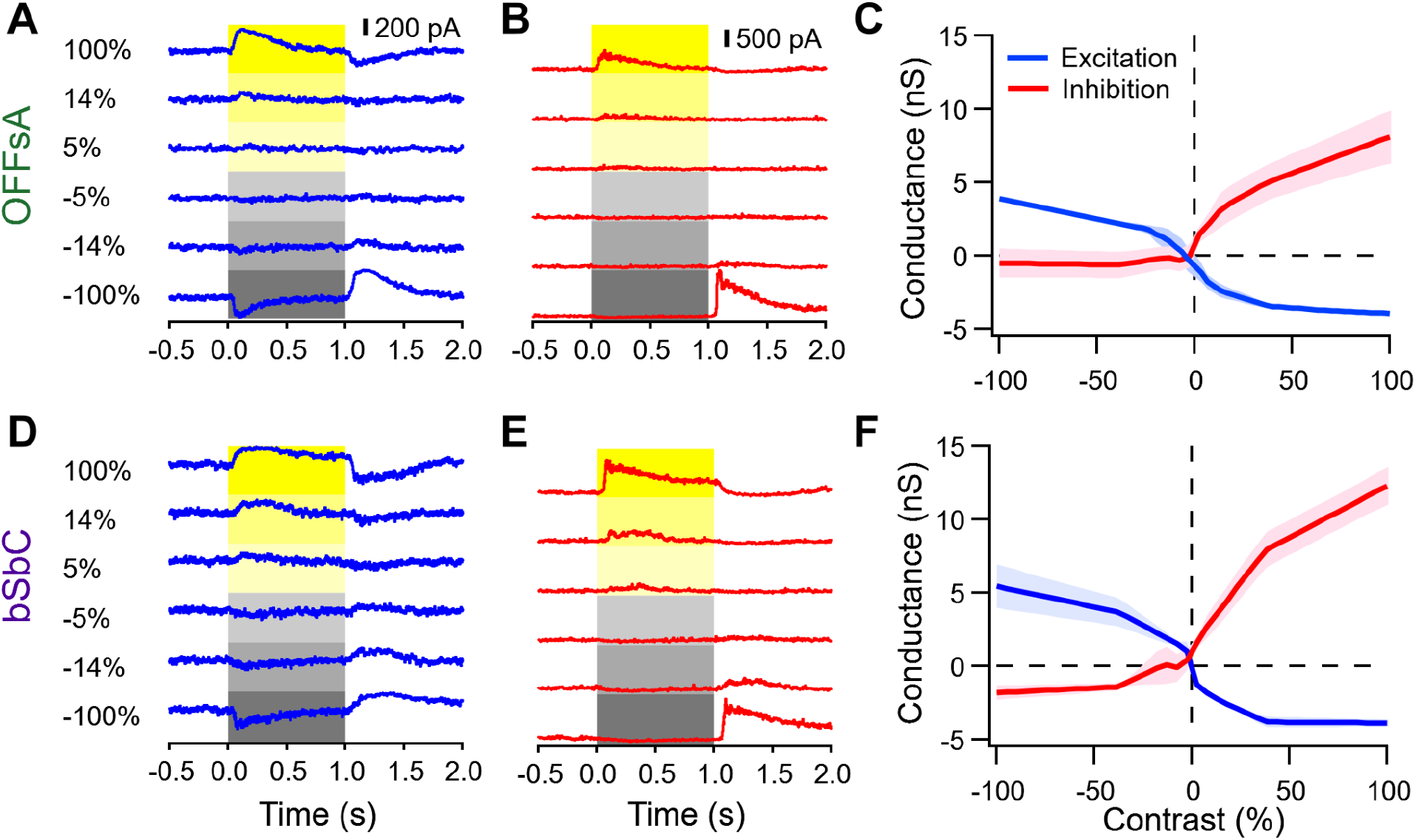
Synaptic currents in OFFsA and the bSbC RGCs are similar. **(A, B, D, E)** Example traces from voltage clamp recordings during the presentation of contrast steps as indicated at a hold of -60 mV to measure excitatory currents **(A, D)** and at a hold of 20 mV to measure inhibitory currents **(B, E)** from an OFFsA RGC **(A, B)** and a bSbC RGC **(D, E)**. **(C, F)** Peak excitatory (*blue*) and inhibitory (*red*) synaptic conductance as a function of contrast for OFFsA RGCs (**C)** and bSbC RGCs (**F)**. Shaded regions are SEM across cells (N = 3 OFFsA, 3 bSbC).

Given that the synaptic conductances appeared so similar, we hypothesized that they might be functionally interchangeable. To test this hypothesis, we injected the synaptic currents measured in one RGC type into the other RGC type and measured spikes. We used dynamic clamp (Sharp et al., 1993) to inject the measured synaptic inputs, recorded in response to 100% negative and positive contrast steps, into both RGC types as conductances (**Figure 4**). By permuting the identities of the synaptic conductances with respect to the identities of the RGCs, we were able to isolate the influence of synaptic input from that of intrinsic properties. We first determined the identity of OFFsA and bSbC RGCs with cell-attached recordings of light responses. Then, we blocked their synaptic input with a pharmacological cocktail (see **Methods**) and simulated synaptic input with currents injected through a whole-cell electrode. In the ‘matched’ condition, we injected excitatory and inhibitory conductances matched to the RGC type in which they were recorded (**Figure 4A**, traces **i** and **iii**). In the ‘swapped’ condition, we injected synaptic conductances measured in OFFsA RGCs into bSbC RGCs and vice versa (**Figure 4A**, traces **ii** and **iv**).

**Figure 4.**
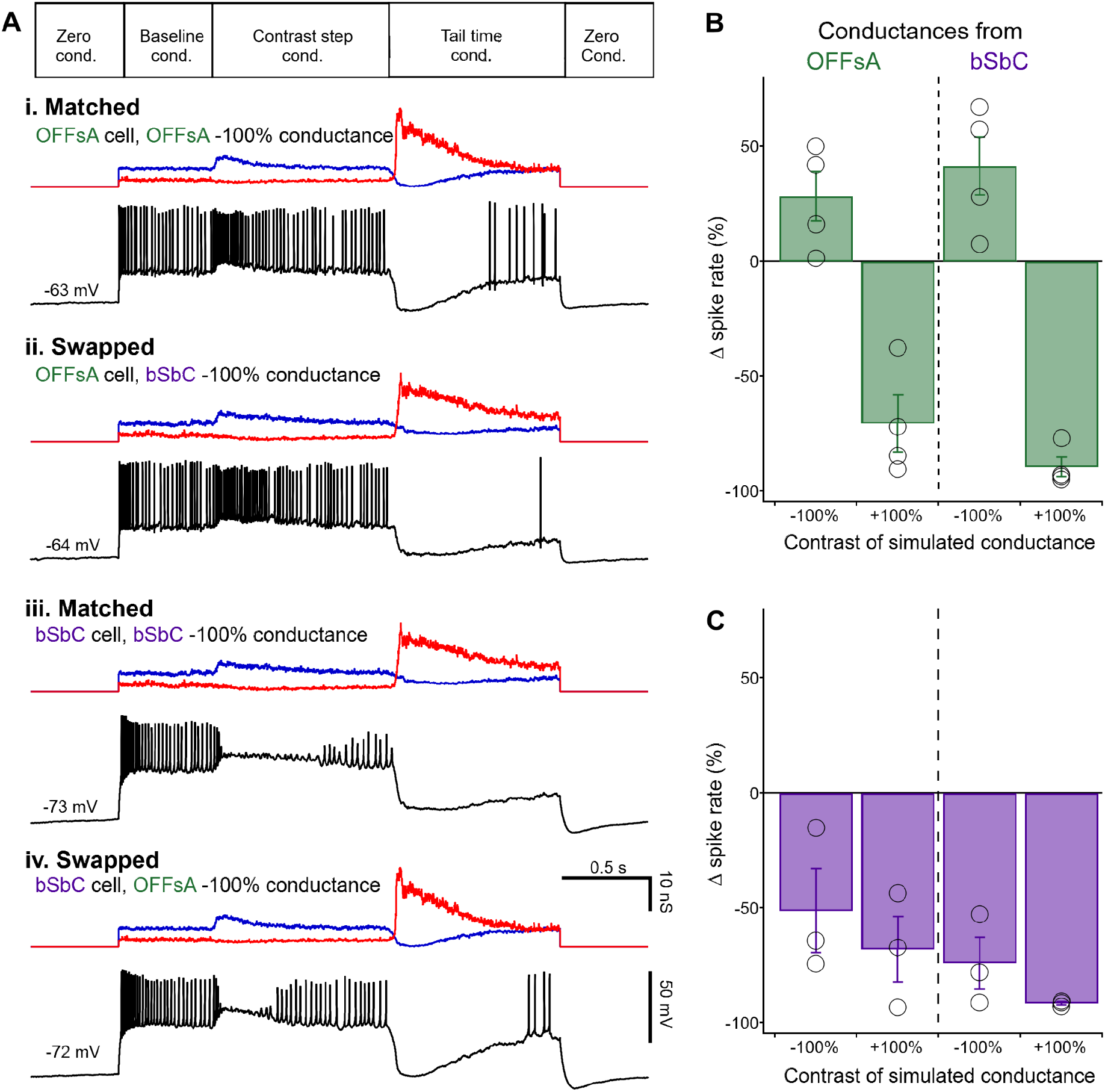
Synaptic conductances in OFFsA and bSbC are functionally interchangeable. **(A)** Dynamic clamp protocol (t*op*) and example current traces (*black*) from 4 conditions as indicated. Excitatory (*blue*) and inhibitory (*red*) input conductances are shown above each trace. **(B)** Change in spike rate during the first 0.5 s after simulated stimulus onset measured in OFFsA RGCs for simulated OFFsA conductances (*left*) and bSbC conductances (*right*) for both negative and positive contrast conditions. **(C)** Same as **(B)** for bSbC RGCs. OFFsA N = 4, bSbC N = 3.

For both OFFsA and the bSbC RGCs, the spike pattern for simulated positive and negative contrast aligned with the identity of the recorded cell rather than the identity of the injected conductances (**Figure 4B**,**C**). That is, OFFsA RGCs increased firing for simulated negative contrast conductances regardless of whether those conductances had been measured in OFFsA or bSbC RGCs, and bSbC RGCs went into depolarization block for simulated negative contrast conductances regardless of the cell type in which they had been measured.

Dynamic clamp is an imperfect approximation of the pattern of synaptic input that cells receive in natural conditions, in part because currents are delivered to the soma rather than the dendrites. In practice, experimentalists scale the recorded conductances down when they are injected in order to elicit spike patterns that resemble those measured with natural inputs. Large discrepancies in these scale factors between cells would suggest important differences in dendritic processing or input resistance that failed to be recapitulated in our experiments. On the contrary, we found no significant difference in the scale factors between the two cell types (**Figure S1A**, p = 0.70). In addition, we found no significant difference between the baseline firing rate found in whole-cell current clamp recordings and the simulated baseline in our dynamic clamp recordings in either cell type (**Figure S1B**; OFFsAs, N = 6, 4, p = 0.67; bSbCs, N = 6, 3, p = 0.45). Thus, our dynamic clamp experiments support the conclusion that the opposite responses of OFFsA and bSbC RGCs to negative contrast stimuli arise primarily from differences in the intrinsic properties that contribute to depolarization block.

### Spike waveforms differ between OFFsA and bSbC RGCs

Increased susceptibility to depolarization block caused bSbC RGCs to stop firing action potentials for the same inputs that caused OFFsA RGCs to fire at high rates (**Figure 4**). We performed whole-cell current clamp recordings in the dark to examine different physiological properties that could contribute to differential responses in the two cell types (**Figure 5A**). We then measured two passive electrical properties of OFFsA and bSbC RGCs that could contribute to differences in their susceptibility to depolarization block: input resistance and resting membrane potential (**Figure 5B**,**C**). Neither parameter differed significantly between cell types (input resistance, p = 0.15, resting potential, p = 0.12; N = 10 OFFsA, 9 bSbC), but they both trended towards the bSbC RGC being more excitable than the OFFsA. Despite their similar membrane properties, OFFsA and bSbC RGCs had different spontaneous spike waveforms in darkness (**Figure 5D-F**). Both spike peak and maximum slope were significantly smaller in bSbC RGCs than in OFFsA RGCs (spike peak: -5.0 ± 1.2 mV vs. 0.6 ± 1.4 mV, p = 0.0068; max slope: 89 ± 9 V/s vs. 180 ± 30 V/s, p = 0.012, N = 9, 10).

**Figure 5:**
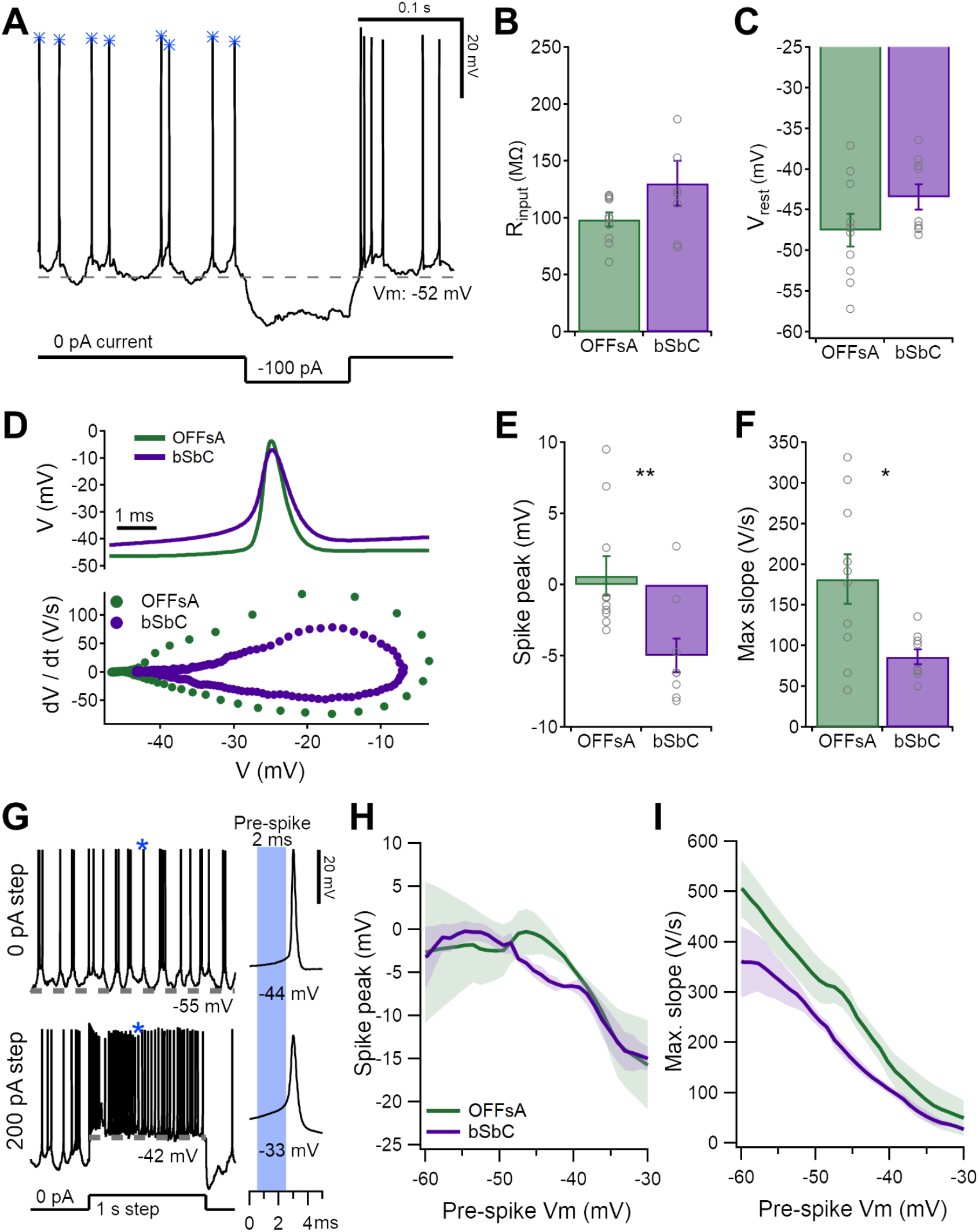
Spike waveforms of the OFFsA and bSbC reveal differences in active conductances. **(A)** Raw data trace from a cell recorded at rest, in darkness with spontaneous spikes. A hyperpolarizing step was used to calculate input resistance **(B)**. *Gray* dotted line denotes the resting membrane potential **(C)**. The *blue* asterisks show which spikes were included for analysis in **(D-E)**. **(B, C)** Passive electrical properties of OFFsA and bSbC RGCs: input resistance (OFFsA N = 10, bSbC N=9, p = 0.13) **(B)** and resting membrane potential (p = 0.16) **(C)**. **(D)** Representative average spike waveforms for OFFsA (*green*) and bSbC (*purple*) RGCs (*top*) and their accompanying dV/dt plots (*bottom*). **(E, F)** Parameters of spontaneous spike waveforms in OFFsA and bSbC RGCs: spike peak (p = 0.0068) **(E)** and maximum rising slope (p = 0.013) **(F)**. ** p < .01; * p < .05. **(G)** Example traces from a cell at rest with 0 pA of baseline current with current steps of 0 pA (*top*) and 200 pA (*bottom*). A spike of interest is indicated with a *blue* asterisk and is shown expanded at right. The pre-spike Vm was calculated using the 2 ms prior to threshold (*blue* shaded region). **(H, I)** Spike peak **(H)** and maximum rising slope **(I)** as a function of the voltage immediately preceding each spike. Shaded region is SEM across cells (N = 3 for each RGC type).

To isolate differences in spike generation between OFFsA and bSbC RGCs we elicited spikes with depolarizing current injections from a range of different baseline voltages (**Figure 5G**). As expected from sodium channel inactivation, for both RGC types, spikes elicited from more depolarized potentials were smaller (**Figure 5H**) and had shallower slopes (**Figure 5I**). Maximum action potential slope is proportional to sodium current multiplied by membrane capacitance (Hille, 2001; Hodgkin and Huxley, 1952). Membrane capacitance was similar between the two RGC types (p = 0.45, see **Table S1**), thus, the fact that bSbC RGCs had spikes with smaller slopes than those OFFsA RGCs across voltages indicated that the total sodium conductance in the bSbC RGCs was lower than that in OFFsA RGCs. This difference in total sodium conductance could arise from differences in sodium channel densities or differences in channel properties, and we explored both of these possibilities in experiments described below.

### Small and shallow somatic spikes are not propagated down the axon

Action potentials only transmit visual information from the retina to the brain if they are propagated down RGC axons. The OFFsA and bSbC RGCs have differences in the maximum slopes of their spike waveforms that persist across a range of physiological membrane voltages (**Figure 5**). Spike peak and maximum slope both impact the probability of a spike propagating down an axon (Khaliq and Raman, 2005; Milner and Do, 2017), and we questioned whether the two cell types had differential spike propagation. To answer this question, we made whole-cell recordings from the soma while performing axon-attached recordings from blebs of the same cell’s axon (at distances from 45 μm to 474 μm) (**Figure 6A**). We then used depolarizing current injections of different amplitudes to cause sodium channel inactivation, and we measured whether each spike successfully propagated down the axon (**Figure 6B**). All cells had at least 2000 somatic spikes that we analyzed to generate curves of propagation likelihood as a function of spike peak (**Figure 6C)** and maximum slope (**Figure 6D**). We performed the experiment in two RGCs of each type. The spike peak voltage required for 50% propagation (V_1/2_) in OFFsA RGCs was -17.2 mV and -19.7 mV; V_1/2_ in the OFFsA RGCs was -21.4 mV and -14.8 mV. Maximum slope values required for 50% propagation (V/s_1/2_) were also similar in the two RGC types (15.9 V/s and 43.7 V/s OFFsA RGCs vs. 21.2 V/s and 26.1 V/s in bSbC RGCs). We concluded from these data that, as in cerebellar Purkinje cells (Khaliq and Raman, 2005), maximum slope is a good predictor of the probability of an action potential propagating down the axon and that the two RGC types do not differ systematically in the relationship between maximum slope and propagation likelihood. We applied a criterion maximum slope of 20 V/s to spike waveforms when counting action potentials from intracellular recordings in either RGC type (used in **Figure 2C** and throughout). Therefore, the differences we measured in the spike waveforms of these two RGC types (**Figure 5**) would be predicted to translate to differences in visual information about contrast sent to the brain.

**Figure 6.**
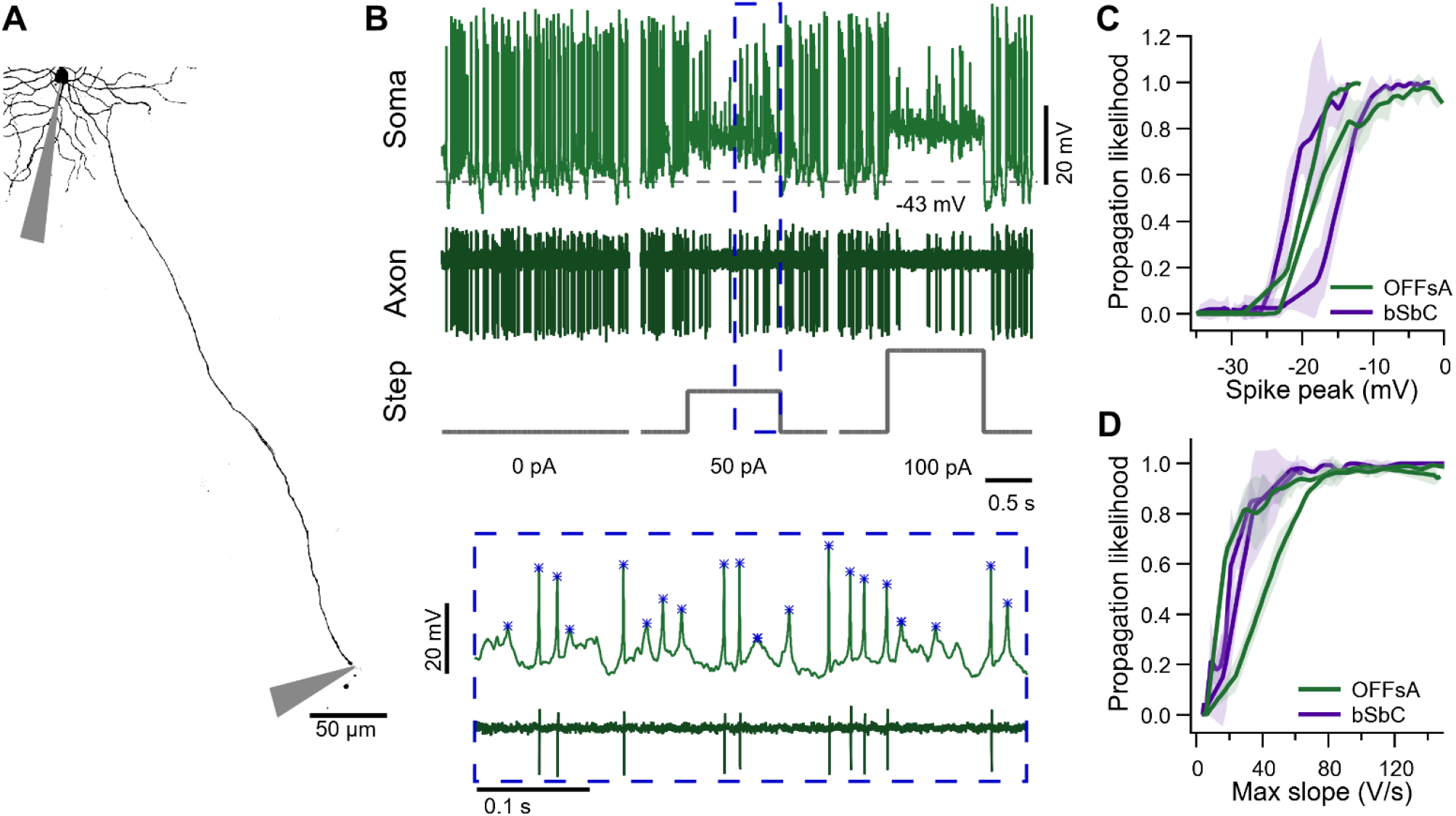
Spike propagation depends on spike waveforms similarly in OFFsA and bSbC RGCs. **(A)** A Z-projected image of an OFFsA RGC filled with Alexa 488 showing the positions of the somatic and axonal electrodes. **(B)** Traces from the soma (*top*) and axon (*bottom*) electrodes for the cell in **(A)** along with injected current steps (*bottom)*. Inset shows magnified view with analyzed somatic spike events marked with asterisks. **(C)** Spike propagation likelihood as a function of the peak of the somatic spike in OFFsA (*green*) and bSbC (*purple*) RGCs. Shaded regions are 2 standard deviations from the mean over randomly subsampled groups (see **Methods**). **(D)** Same as **(C)** but as a function of the maximum rising slope of the somatic spike. Axons were recorded at distances of 210 μm and 252 μm (OFFsA); 45 μm and 474 μm (bSbC).

### OFFsA and bSbC RGCs have differences in sodium channel types and AIS length

We revealed that the bSbC RGCs consistently had shallower spikes than the OFFsA across voltages (**Figure 5**) and that these differences translate to the likelihood that a spike will propagate (**Figure 6**). The maximum slope differences observed indicate that the total sodium conductance is different in the two cell types. We wanted to test if differences in sodium channel subtypes could contribute. The two primary sodium channel subtypes in the adult mouse retina are Na_V_1.2 and Na_V_1.6 (Van Hook et al., 2019). Na_V_1.6 channels are the canonical sodium channels at the AIS in most spiking neurons, and they are the lowest voltage activated among Na_V_ channels (Catterall et al., 2005; Llinas, 1988; Royeck et al., 2008; Rush et al., 2005). These channels are also principal carriers of resurgent current that is important for maintaining high spike rates in some neurons, like Purkinje cells (Raman et al., 1997). In RGCs, Na_V_1.6 channels have been found in the AIS of every RGC type in which they have been studied (Raghuram et al., 2019; Van Hook et al., 2019). We used 4,9-anhydrotetrodotoxin (49TTX), a Na_V_1.6-specific channel blocker, to examine if the two cell types express different functional subtypes of sodium currents (Browne et al., 2017; Rosker et al., 2007). We chose a concentration of 49TTX (10 nM) near the reported IC_50_ for Na_V_1.6 channels (7.8 nM) and more than 30 times below the next lowest IC_50_ for a different Na_V_ channel type (341 nM) so that we could be certain not to observe off-target effects at the expense of an incomplete block of Na_V_1.6 (Rosker et al., 2007).

We applied 49TTX to the bath (in the presence of synaptic blockers) while recording current-induced spikes in OFFsA and bSbC RGCs (**Figure 7**). In OFFsA RGCs, blockade of Na_V_1.6 channels affected spike waveforms in the expected ways; they became smaller, and shallower (**Figure 7A**,**C**,**D**). However, in bSbC RGCs, we observed no effect in spike peak or maximum slope across the physiological range of pre-spike voltage (**Figure 7B**,**C**,**D**). There was no effect on either input resistance (**Figure S2A**, OFFsA p = 0.83, and bSbC p = 0.28, or on resting membrane potential (**Figure S2B**, OFFsA p = 0.29, and bSbC p = 0.91). Thus, the sodium currents in the OFFsA are sensitive to 49TTX, while the bSbC sodium currents are resistant to block by 49TTX. These results suggest that unlike in all other RGCs in which sodium channel composition has been measured (Van Hook et al., 2019), spike generation in bSbC RGCs is not driven by Na_V_1.6 channels.

**Figure 7.**
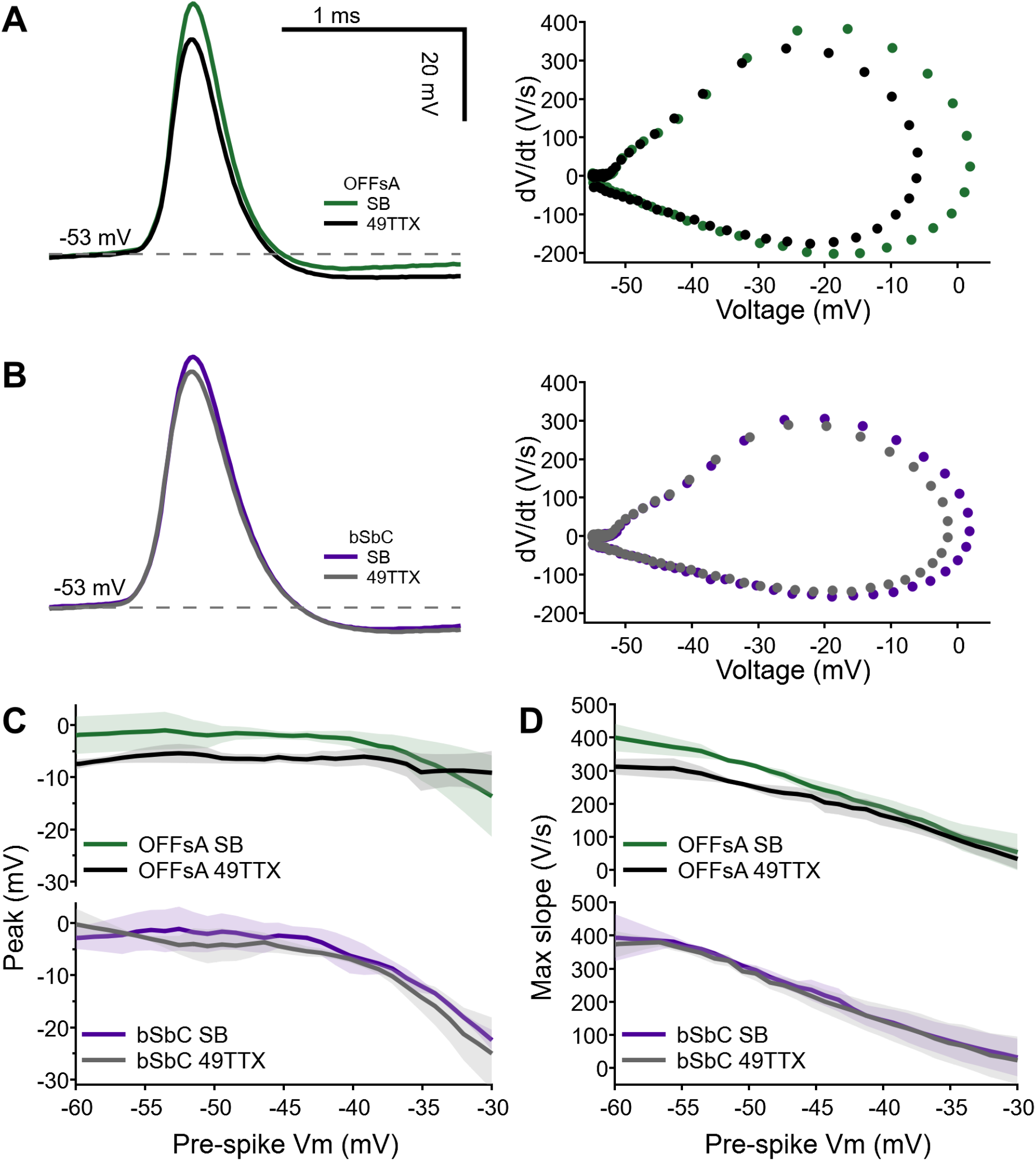
An Nav1.6 specific channel blocker affects the OFFsA and not the bSbC. **(A, B)** Example action potentials waveforms and corresponding dV/dt plots in synaptic blocker (SB) conditions and with 10 nM 49TTX for the OFFsA **(A)** and bSbC **(B)**. OFFsA is green for SB and black for 49TTX, bSbC is purple for SB and gray for 49TTX. **(C, D)** Plots of action potential peak **(C)** and max slope **(D)** over the local membrane voltage. Shaded region is SEM across cells (see **Methods**, N=3 for both types).

To investigate possible differences in Na_V_ channel densities between OFFsA and bSbC RGCs, we imaged the AIS, where Na_V_ channels, typically Na_V_1.2 and Na_V_1.6 in RGCs (Raghuram et al., 2019; Van Hook et al., 2019), are tethered at high density with Ankyrin G (AnkG) (Bender and Trussell, 2012; Boiko et al., 2003). We used an antibody against AnkG in OFFsA and bSbC RGCs that had been filled with Neurobiotin via patch electrodes (**Figure 8**) and used a threshold on the Neurobiotin-masked AnkG signal to determine the start and end of the AIS (see **Methods**). Consistent with previous reports of cortical neurons and RGCs (Hamada et al., 2016; Höfflin et al., 2017; Raghuram et al., 2019), we found bSbC RGCs with both axo-somatic (4/6) and axo-dendritic morphology (2/6). We found only axo-somatic OFFsA RGCs (N = 4). We compared AIS parameters in the axo-somatic population of OFFsA RGCs to those in bSbC RGCs. The diameter of the AIS was similar in the two RGC types (1.39 ± .074 µm vs. 1.35 ± .039 µm, p = 0.66, **Figure 8C**) as was the distance from the soma (23 ± 4.9 µm in OFFsA vs. 19 ± 4.4 µm in bSbC, p = 0.22, **Figure 8D**), but the AIS was shorter in bSbC RGCs than in OFFsA RGCs (15 ± 3.4 µm vs. 21 ± 2.5 µm, p = 0.03, **Figure 8E**). If the density of Na_V_ channels in the AIS is similar between these RGCs, the shorter AIS in bSbC RGCs (29% shorter than in OFFsA RGCs) would account for a substantial fraction of the computed difference in maximum sodium conductance (37% lower in bSbC RGCs than OFFsA RGCs, **Table S1**).

**Figure 8:**
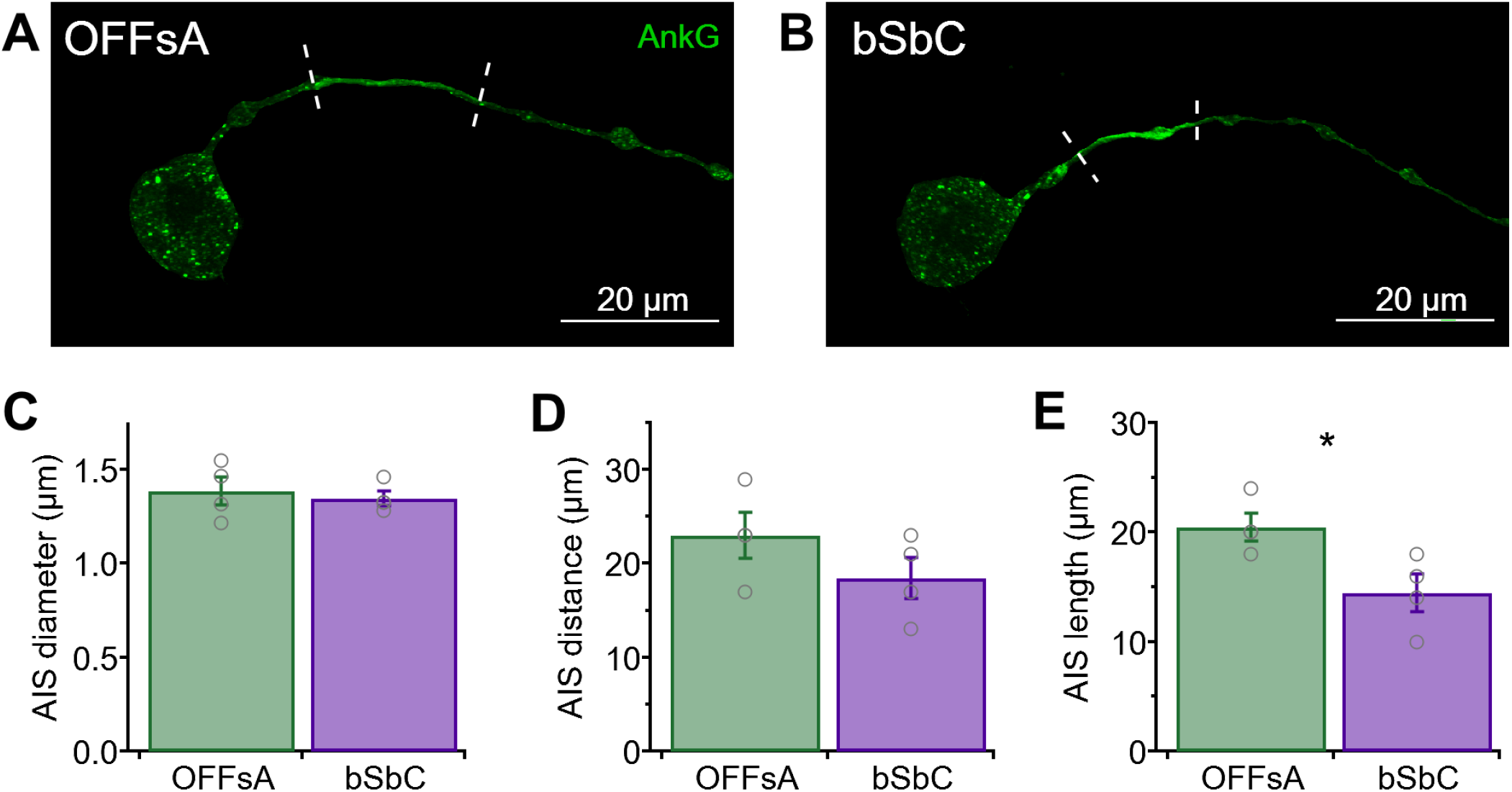
The OFFsA and bSbC have differences in AIS structure. **(A, B)** Ankyrin G staining masked by the cell fills in an OFFsA RGC **(A)** and a bSbC RGC **(B)**. Dashed lines indicate the boundary of the AIS as determined by thresholding. **(C-E)** AIS diameter (p = 0.66) **(C)**, distance (p = 0.22) **(D)**, and length (p = 0.03) **(E)** for OFFsA (N= 4) and bSbC (N = 4) RGCs. * p < 0.05.

### A compartmental model of spike generation in OFFsA and bSbC RGCs

To establish a quantitative framework for our understanding of how each of the properties we measured in OFFsA and bSbC RGCs contributes to their different susceptibility to depolarization block, we built a compartmental model of spike generation in these RGCs. We modeled the dendritic tree as a single compartment and included separate compartments for the soma, axon hillock, AIS, and axon (**Figure 9A**). Model parameters were calculated from our electrophysiological measurements or taken from literature values (**Table S1**, see **Methods**).

**Figure 9:**
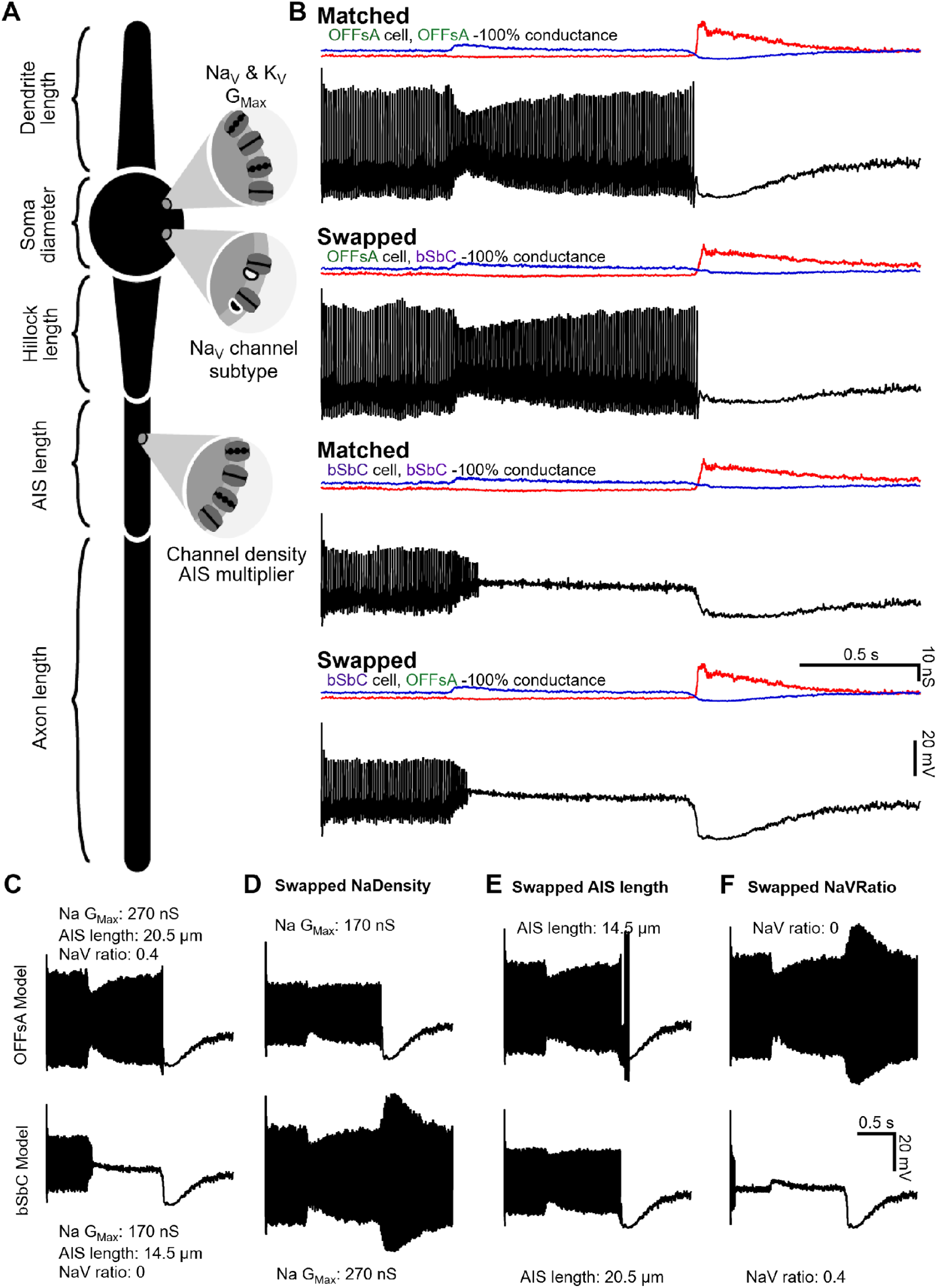
Modeling demonstrates the interplay of sodium channel properties with morphological properties and their effect on cell excitability. **(A)** Model schematic labeling most of the key parameters. Values for all parameters can be found in **Table 1**. **(B)** OFFsA and bSbC models that only differ in their AIS length (OFFsA = 20.5 μm, bSbC 15.5 μm), ratio of Na_V_1.6 (OFFsA = 40%, bSbC = 0%), and the total sodium conductance of the cell (OFFsA = 272.8 nS, bSbC = 169.4 nS). Then each cell was injected with the conductances measured in **Figure 4** and the spiking outputs are shown in *black*. Excitatory (blue) and inhibitory (*red*) input conductances are shown above each trace. **(C-F)** Traces for the OFFsA model (top row) and bSbC model (bottom row) in response to current injections. In **(D-F)** the parameters for sodium channel total conductance for the cell **(D)**, AIS length **(E)**, and the ratio of Na_V_1.6 channels **(F)** were swapped between the OFFsA and bSbC models.

All models include simplifying assumptions, and ours is no exception. The complement of active channels in our models was limited to the fast Na_V_ channels and a generic inward rectifying K_v_ channel despite observations that some RGC somas also include HCN channels, a slow voltage-gated sodium channel (Na_V_1.8), several types of Ca^2+^ channels, and/or different types of K_V_ channels (Van Hook et al., 2019). Our model also excluded dendritic processing as an intrinsic property that contributes to cell-type specific computations in RGCs (Fohlmeister and Miller, 1997) to focus on the spike-generation differences we measured with somatic dynamic clamp (**Figure 4**). Models for Na_V_1.2 and Na_V_1.6 sodium channels were taken from somatic and axonal outside-out patch recordings in rat pyramidal neurons (Hu et al., 2009). The two models differed in the voltage dependence of activation where the half-activation voltage (V_1/2_) for Na_V_1.2 was -43 mV and the slope was 7 mV whereas for Na_v_ the values were -42 mV and 6 mV respectively. There was no difference in the voltage dependence of inactivation included in the models. The Na_V_1.6 model did not incorporate the resurgent current that is known to play a role in high frequency spiking in some neurons (Khaliq et al., 2003). Voltage gated potassium channels are an even more diverse class than Na_V_ channels (Hille, 2001), and they could also contribute to differences in susceptibility to depolarization block between bSbC and OFFsA RGCs. Our decision to leave K^+^ channel diversity out of our model was influenced by the fact that we observed significant differences in the rising phase of action potentials (**Figures 5**,**7**) which is controlled primarily by Na_V_ channels (Hodgkin and Huxley, 1952). Therefore, while our model did not capture all aspects of intrinsic properties of RGCs, it provided a useful tool for studying the contributions of components that are not easily isolated experimentally.

First, to see if the measured intrinsic differences between OFFsA and bSbC RGCs were sufficient to reproduce their different responses to the same synaptic currents, we fixed all parameters of the model except for three – total Na_V_ conductance, Na_V_1.6 fraction (with the remainder Na_V_1.2), and AIS length – and stimulated the models with the recorded conductances as in our dynamic clamp experiments (**Figure 4**). Values for the three parameters that differed between the OFFsA and bSbC RGC models were calculated directly from our measurements in the two cell types (see **Methods**). Indeed, the different values of these three parameters were sufficient to produce the pattern of results we observed experimentally: sustained spiking in the OFFsA RGC model and depolarization block in bSbC RGC model for identical input currents (**Figure 9B**).

To study the role of each parameter individually, we fixed the synaptic conductances to those for the OFFsA and swapped one model parameter between the two cells at a time so that we could assess each parameter’s contribution to susceptibility to depolarization block. These changes did not affect the total resistance of the models appreciably (**Table S2**). The OFFsA model did not undergo depolarization block when we swapped in any one the individual bSbC parameters: total conductance (**Figure 9D**, top trace), AIS length (**Figure 9E**, top trace), or Na_V_1.6 ratio (**Figure 9F**, top trace). The bSbC model was rescued from depolarization block when we swapped in the OFFsA value for either the total sodium channel conductance (**Figure 9D**, bottom trace) or AIS length (**Figure 9E**, bottom trace). When we swapped the OFFsA value for Na_V_1.6 ratio (i.e. added Na_V_1.6 channels) into the bSbC model (**Figure 9F**, bottom trace), the model prematurely went into depolarization block at the start of the conductance trace. This behavior is explained by the fact that a larger proportion of Na_V_ channels was inactivated at rest (−45 mV) in this model due to the lower voltage activation of Na_V_1.6.

Together, our models suggest that the high susceptibility to depolarization block in bSbC RGCs results from a coordination of all three of these intrinsic properties of their spike generator. Low total Na_V_ conductance and a short AIS conspire to create a smaller pool of sodium channels available for spike generation, while the absence of Na_V_1.6 allows these cells a slightly larger resting potential range in which they remain unblocked so that they can spike in baseline conditions but block for small depolarizations.

## Discussion

RGCs respond to particular visual features, and their selectivity is typically explained by the excitatory and inhibitory synaptic inputs they receive from the complex network of upstream interneurons in the retina: bipolar and amacrine cells. Most models of RGC feature selectivity consider spike generation as an afterthought; simple thresholds or leaky-integrate-and-fire models have been the standard way to translate feature-selective inputs into spikes (Joesch and Meister, 2016; Johnson et al., 2018; Nath and Schwartz, 2017; Venkataramani and Taylor, 2016; Venkataramani et al., 2014; Zhang et al., 2012). The diversity of spike patterns in RGCs to the same somatic current injections was an indication that spike generation could play a role in RGC feature selectivity (O’Brien et al., 2002; Wong et al., 2012). This role has been confirmed in recent work on ipRGCs (Emanuel et al., 2017; Milner and Do, 2017; Sonoda et al., 2018), but it was not known whether and how properties of spike generation affect feature selectivity in other RGC types.

This study augments our understanding of the role of spike generation in RGC feature selectivity by introducing an RGC type, the bSbC RGC (**Figure 1**), whose high susceptibility to depolarization block creates a situation where its firing rate *decreases* for negative contrast despite the depolarizing influence of a strong excitatory current and a reduction of inhibition (**Figures 2**,**3**). We compared the bSbC RGC to the well-studied OFFsA RGC and showed that, despite functionally interchangeable synaptic inputs (**Figure 4**), these RGCs have dramatically different contrast response profiles in their spike output (**Figure 2**). OFFsA and bSbC RGCs also differ in their spontaneous spike waveforms (**Figure 5**) suggesting differences in Na_V_ conductances. The two cell types do not differ in how they propagate spikes down the axon (**Figure 6**), thus the spiking differences we measured at the soma lead to differences in contrast information sent to the brain. We used pharmacology (**Figure 7**), imaging of the AIS (**Figure 8**), and compartment modeling (**Figure 9**) to implicate three biophysical differences between these RGC types that drive this functional difference in spike generation: total sodium channel conductance, AIS length, and the complement of sodium channel subtypes.

### Functional and mechanistic differences among SbC RGCs

Suppressed-by-contrast (SbC) RGCs are defined by their reduction of tonic firing for both positive and negative contrasts. The SbC group comprises multiple distinct RGC types (Jacoby and Schwartz, 2018; Mastronarde, 1985) and represents a substantial fraction (15–25%) of the retinal input to the dorsal lateral geniculate nucleus of the thalamus (Liang et al., 2018; Piscopo et al., 2013; Román Rosón et al., 2019). The two SbC RGC types previously identified in mice share a circuit motif in which ON and OFF inhibition dominates a much smaller excitatory drive (Jacoby et al., 2015; Mani and Schwartz, 2017; Tien et al., 2015, 2016), but they differ in the details of their upstream circuits and in the spatial and temporal tuning of spike suppression for negative versus positive contrasts (Jacoby and Schwartz, 2018). The bSbC RGC represents a third SbC type in the mouse, and it has a striking asymmetry in its contrast-suppression mechanism; positive contrast leads to spike suppression by hyperpolarization, while negative contrast leads to spike suppression by depolarization. The duration of spike suppression in bSbC RGCs depends on the amplitude of the contrast signal for both contrast polarities, but it is longer for high positive than high negative contrasts (**Figure S3**). Future work will make more comprehensive functional comparisons between the three bSbC RGC types in mice (and any more that are discovered) to provide insights into the possible behavioral niche of each type and how each is supported by a different circuit or cell-intrinsic mechanism.

### Depolarization block as a mechanism for neural computation

We show that bSbC RGCs undergo depolarization block in response to negative contrast stimuli, suggesting that it is a natural physiological state rather than a pathological state. Depolarization block clearly can be a pathological state at the cellular level, and it has been associated with pathological brain states like epilepsy (Bikson et al., 2003). For more than three decades, there has been evidence that dopamine neurons undergo depolarization block in response to treatment with antipsychotics without clear detrimental effects to their long-term survival (Grace, 1992; Valenti et al., 2011), but this is still a state produced by an artificial drug. A number of groups have argued that depolarization block can be a naturally occurring state *in vivo* in the hippocampus (Bianchi et al., 2012; Bragin et al., 1997; Knauer and Yoshida, 2019) but none have recorded it directly. Perhaps the best evidence of depolarization block being used for neural coding comes from sensory neurons. Olfactory sensory neurons in mice use this state in an *ex vivo* preparation of the olfactory epithelium (Ghatpande and Reisert, 2011), and the previous paper on depolarization block in M1 ipRGCs included indirect evidence that the cells behave similarly *in vivo* (Milner and Do, 2017). Neurons in the suprachiasmatic nucleus downstream of M1 ipRGCs have also been shown to undergo depolarization block across the day-night cycle (Belle et al., 2009). Our work provides further support for the view that depolarization block can be a physiological state important for neural coding. In bSbC RGCs, this state would be expected to be quite frequent in natural environments, occurring whenever the cell’s receptive field encounters negative contrast. The additional biophysical and metabolic specializations (e.g. ion pumps) that make depolarization block a sustainable state in bSbC RGCs are potential avenues for future research that could offer insights into how other neurons survive frequent, sustained depolarization.

### Spike waveforms may be useful for electrically fingerprinting RGCs in functionally damaged retina

Retinal prostheses require models to translate incoming light patterns into electrical patterns on the implanted electrode array, and these models depend critically on the correct identification of RGC types (Fried and Werblin, 2006; Werginz et al., 2020). For these models to be successful in restoring sight to blind patients, they must work in retinas where light responses are largely or completely absent. In these conditions, spike waveforms and spontaneous spike train statistics are some of the only available information about RGC typology (O’Brien et al., 2002; Wong et al., 2012; Zeck and Masland, 2007). As has been shown for the alpha RGCs, spike waveforms can help distinguish RGC types, even in extracellular recordings (Krieger et al., 2017). Our work adds a mechanistic understanding to the spike waveform differences between OFFsA and bSbC RGCs, adding to the foundation of work on electrical fingerprinting of RGCs.

## Acknowledgements

We are thankful to all Schwartz Lab members for their feedback and technical assistance through the project. Thanks to Devon Greer for designing and making the schematic of the model. We would like to acknowledge Indira Raman, Tiffany Schmidt, Steven DeVries, Soile Nymark, Michael Tri Do, Julia Fadjukov, and Zachary Jessen for their feedback and comments on the manuscript. Funding for this research was provided by National Institutes of Health Grant F31 EY030737.

## Author Contributions

S.W. and G.W.S. conceived the study plan and designed the experiments. S.W. and G.W.S. collected and analyzed the data. S.W. and G.W.S. wrote the paper.

## Methods

### Animals

Wild-type mice on the C57Bl/6 background of either sex and at least 4 weeks of age were used for all experiments. Animals were used in concordance with protocols approved by Northwestern University Institutional Animal Care and Use Committee.

### Electrophysiology

Retinas were dissected in infrared light then perfused with oxygenated Ames medium at 31-32 °C at a rate of 10 mL/min (Jacoby et al., 2015; Nath and Schwartz, 2016).

#### Typology

RGC types were established by performing cell-attachedd recordings to both light steps (200 μm diameter, 200 R*/rod/s from darkness) and contrast response stimuli (5 logarithmically spaced steps from 2% to 100% contrast both positive and negative, from a background of 1000 R*/rod/s). Stimuli were presented for 1 s each with the blue LED (450 nm) on a digital projector (LightCrafter 4500, Texas Instruments). Glass electrodes were pulled to 2-3 MΩ and filled with Ames medium. All loose patch recordings were collected at a sample rate of 10 kHz. Details of our criteria for functional classification of RGCs can be found in (Goetz et al.).

#### Whole-cell recordings

Voltage clamp recordings were performed with a cesium based internal solution (104.7 mM cesium methanesulfonate, 10 mM TEA-Cl, 20 mM HEPES, 10 mM EGTA, 2 mM QX-314, 5 mM ATP, 0.5 mM GTP, cesium hydroxide to adjust pH to approximately 7.2). To measure excitatory synaptic currents the holding potential was -60 mV, and for inhibitory currents the holding potential was 20 mV (Nath and Schwartz, 2016). Whole cell voltage clamp recordings were recorded at 10 kHz. Current clamp recordings were performed with a potassium aspartate-based (K-aspartate) solution (125 mM L-Aspartic Acid Potassium Salt, 1 mM MgCl, 10 mM KCl, 10 mM HEPES, 2 mM EGTA, 1 mM CaCl_2_, 4 mM ATP, 0.5 mM GTP, KOH to adjust pH to approximately 7.2). Whole-cell recordings used an electrode of 4-6 MΩ. Current clamp recordings were collected at a sampling rate of 50 kHz. All recordings were obtained using a 2-channel patch-clamp amplifier (Multiclamp 700B, Molecular Devices).

#### Analysis

For whole-cell current-clamp recordings, spikes were detected using an algorithm that finds local maxima with a prominence of at least 5 mV. Based upon the failure rate analysis (see below, Axon Recordings), spikes with a maximum slope less than 20 V/s were filtered out and not considered in any following analyses. The baseline membrane potential (V_m_) of the epoch and input resistance were calculated from low pass filtering the data to remove spikes using a moving median filter set at 101 samples. Action potential waveforms were then analyzed using custom MATLAB code. Threshold was defined as the place where the second derivative of the voltage trace exceeded 5 times its variance. The pre-spike V_m_ was calculated from the mean 2 ms prior to threshold. Once all the parameters were collected, the data were collected into 1 mV bins. Missing data in the range of -60 mV to -30 mV were linearly interpolated or extrapolated. The data were then averaged across cells, and error bars show standard error of the mean.

### Dynamic clamp

Dynamic clamp recordings were performed in whole-cell current-clamp configuration as described above. Dynamic clamp hardware and software were implemented as described in (Desai et al., 2017). The input conductances were scaled (**Figure S1A**) such that the baseline spiking rate did not significantly differ between cells recorded in current clamp and dynamic clamp (**Figure S1B**). Spike rates for dynamic clamp were calculated from the last 250 ms of the injected baseline conductance (**Figure 5A** schematic) and from the first 500 ms of conductance during the simulated contrast spot.

### Axon recordings

Axon recordings started by patching a cell with a K-aspartate internal solution containing AlexaFluor 488 after performing loose patch recordings for typology. The electrode was then removed from the cell and the axon was imaged using a 2-photon laser (980 nm). At a sufficient distance away (a range of 45 to 474 μm), a new tear in the inner limiting membrane was made to provide access to the axon. The electrode was then placed under the axon and moved up quickly to tear the axon and form a membrane bleb at its sealed end. Loose patch recordings were then made from the bleb using an electrode of 3-4 MΩ while the soma was patched for a second time and current clamp recordings were performed as described above. Multiple images of both the dendrites and length of the axon were then taken after recording was complete to determine the length of the axon.

Failure analysis was built upon the principle of matched filtering. First, somatic spikes were analyzed as above sans thresholding, and the accompanying segment of the axonal trace was extracted. The mean of the axonal traces for large spikes (greater than 30 mV in amplitude) was normalized to have an integral equal to 1 and used as the template. All axon traces were then convolved with the template to compute their projection values. Projection values for the large spikes were fit to a normal distribution, and values for all projections less than a 2.5% of the mean were deemed to be failures. The spikes were randomly subsampled into 4 distinct groups in order to generate error bars on the propagation likelihoods. The average curve for each cell was then fit to a sigmoid of the form y=1/(1+exp(-(x-V_1/2_)/k)) to find the 50% propagation likelihoods.

### Pharmacology

Pharmacology experiments were conducted during current clamp recordings as described above. All solutions were perfused with the same rate and temperature as above. The synaptic blockers solution included CNQX (50 µM, Tocris) and L-AP4 (20 µM, Tocris) to block all glutamatergic transmission and remove synaptic noise. Then, 4,9-anhydrotetrodotoxin (49TTX, 10 nM, Alomone Labs) was added to the solution and recordings resumed after 5 min. Finally, tetrodotoxin (500 nM, Tocris) was added as a control and eliminated all spikes (data not shown). All recorded cells had datasets for both the synaptic blockers and 49TTX conditions, so Loftus-Masson normalization was performed between the two conditions (Loftus and Masson, 1994).

### Imaging and image analysis

#### Live imaging

After performing whole cell recordings with an internal solution containing AlexaFluor 488, cells were imaged using two-photon microscopy as previously described (Jacoby et al., 2018; Nath and Schwartz, 2016).

#### Stratification

To target cells for immunohistochemistry, after typology, cells were filled with 3% Neurobiotin (Vector Labs) in our K-Aspartate internal form above. Retinas were fixed at room temperature for 15 min in 4% paraformaldehyde (Electron Microscopy Sciences). Then they were blocked at room temperature for 2 hours in 3% Normal Donkey Serum (Jackson Labs) and 0.5% Triton (Sigma) in Phosphate Buffer. Retinas were then incubated with primary antibodies (mouse anti-SMI-32, 1:500; goat anti-ChAT, 1:500) for 5 days at 4 °C. After washing, retinas were incubated with secondary antibodies (568 anti-mouse; 647 anti-goat; streptavidin 488, all at a dilution of 1:500) for 2 days at 4 °C. All secondaries were from Life Technologies and all streptavidin conjugates are from Thermo Scientific. Retinas were then mounted using Vectashield Antifade (Vector Labs) and imaged using a 40x oil immersion objective on a Nikon A1 confocal microscope.

#### Morphology analysis

From both two-photon and confocal images, soma diameter was calculated by tracing an outline of the soma using ‘Freehand Selections’ and solving for diameter in FIJI. Similarly, convex area was measured by drawing a polygon around the tips of the dendrites in a flattened view of the image. Stratification analysis was performed using custom MATLAB software (Nath and Schwartz, 2016) based on a published algorithm (Sümbül et al., 2014) after tracing the dendrites using the SNT plug-in in FIJI (Arshadi et al., 2021).

#### AIS identification

Cells were labeled for immunohistochemistry as above. After fixation and blocking, the retinas were incubated with primary antibodies (mouse anti-Ankyrin G, 1:200) for 5 days at 4 °C. After washing, retinas were incubated with secondary antibodies (488 anti-mouse, 1:200; streptavidin 647, 1:500) for 2 hours at room temperature. Retinas were then mounted using Vectashield Antifade and imaged using a 100x oil immersion objective and a Nikon A1 confocal microscope.

Axon initial segment (AIS) analysis was performed by binning the fluorescence along the length of the axon. First, the axon was traced as above, then the NB image was used to mask the AIS image. Then using custom Matlab software, the image was binned into 4 µm segments and the average fluorescence for each segment was calculated. After normalization, AIS distance was calculated at the point at which a 0.95 threshold was crossed, and length was calculated from that original crossing point until a 0.25 threshold was crossed. Finally, the diameter was calculated by using the average width of the fluorescence across five repeat measurements perpendicular to the AIS.

### Modeling

Modeling was performed using Python 3.7.4 and NEURON 7.7. The model morphology is diagrammed in Figure 9A. Conductances modeled included the built in NEURON passive mechanism and sodium channel and potassium channel models from (Hu et al., 2009). All model parameters and associated methods can be found in **Table S1**.

Membrane specific capacitance was set to 1 µF/cm^2^ and axial resistance was set to 200 Ω·cm (Abbas et al., 2013; Schachter et al., 2010). Morphological parameters were determined from measurements performed from both images (see Morphology section) and capacitance measurements (**Table 1**). Capacitance was computed from the input resistance and the tau of small (<100 pA) hyperpolarizing steps. Capacitance measurements were used to compute the total surface area of the model cell. The surface area was then distributed across the different morphological parameters and the remaining was assigned to the dendritic compartment (Dendritic Length parameter). With the exception of the somatic compartment, all compartments had a diameter of 1 micron. All sections had 15 segments.

We were able to estimate channel conductance from our experimental data. Direct measures of channel conductance across the cell or in a single morphological compartment were unfeasible. Channel conductance was estimated using the maximum and minimum slopes of action potentials for sodium and potassium channels, respectively. Current was calculated as the slope of the action potential times the capacitance of the cell. Then using a holding potential of -60 mV and the reversal potential of our solutions, we could calculate the conductance of the sodium and potassium channels of the entire cell. This assumes that the upstroke and downstroke of the action potential were carried solely by sodium and potassium ions respectively. However, we know that many other channels and conductances play a role in RGCs (Van Hook et al., 2019). Channel density was calculated by dividing the total conductance by the total surface area of the cell. The density was assumed to be uniform across the entire cell except at the AIS where it was assumed to be higher by a factor of 30 (Bender and Trussell, 2012) (**Table S1**). Because the allotment of the total surface area changed as the morphological parameters changed the density was adjusted accordingly, except in the case of AIS length where the model was run with both adjusted and unadjusted densities. Na_V_ ratio is the percentage of Na_V_1.6 sodium channels in the cell and the rest are assumed to be Na_V_1.2. Because the OFFsA had an approximately 20% reduction in its max slope from the application of the IC_50_ dose of 49TTX, the maximum value tested for Na_V_ ratio was 0.4. Unless stated, only one parameter and the corresponding channel densities were changed at a time. These parameters estimated from the current clamp data were sufficient to allow us to investigate the effect of sodium conductance, AIS length, and Na_V_1.6 ratio on the susceptibility of a cell to depolarization block.

The temperature of the simulation environment was set to 32 °C and the time step was 0.01 ms. The reversal potential for sodium, E_Na_, was set to 30 mV, E_K_ was set to -90 mV, and E_leak_ was set to -50 mV. Since we are not accounting for other conductances other than voltage gated sodium, voltage gated potassium, and leak, E_leak_ was set so that the resting membrane potential of the model was similar to what was observed. The initial membrane voltage was set to -55 mV. The leak conductance was adjusted such that the measured input resistance of the average values model was measured at 100 MΩ (**Table 2**) (Freed et al., 1992). Varying amounts of current were injected to simulate the current clamp experiments above and the data were analyzed using the same code.

Contrast responses were simulated by injecting measured conductances into our model cells. The NEURON model environment used a dt of 0.1 ms to match our recorded conductance traces. The traces were fed in using the innate SEClamp method with a reversal potential of 0 mV for excitatory traces, and -70 mV for inhibitory traces accounting for the liquid junction potential. The OFFsA and bSbC models differed in their Na_V_1.6 Ratio, AIS Length, and max Na conductance (see parameters in **Table S1**). The input resistances of each model are listed in **Table S2**.

### Statistics

Data are reported as mean ± SEM. P-values were calculated using a two tailed Student’s T-Tests (paired or unpaired as appropriate) unless specified otherwise.

### Resource availability

#### Lead contact

Further information and requests for resources and reagents should be directed to and will be fulfilled by the lead contact, Greg Schwartz.

### Materials availability

This study did not generate new unique reagents.

### Data and code availability

All data reported in this paper will be shared by the lead contact upon request. All code can be found at https://github.com/SchwartzNU.

Any additional information required to reanalyze the data reported in this paper is available from the lead contact upon request.

## Key resources table

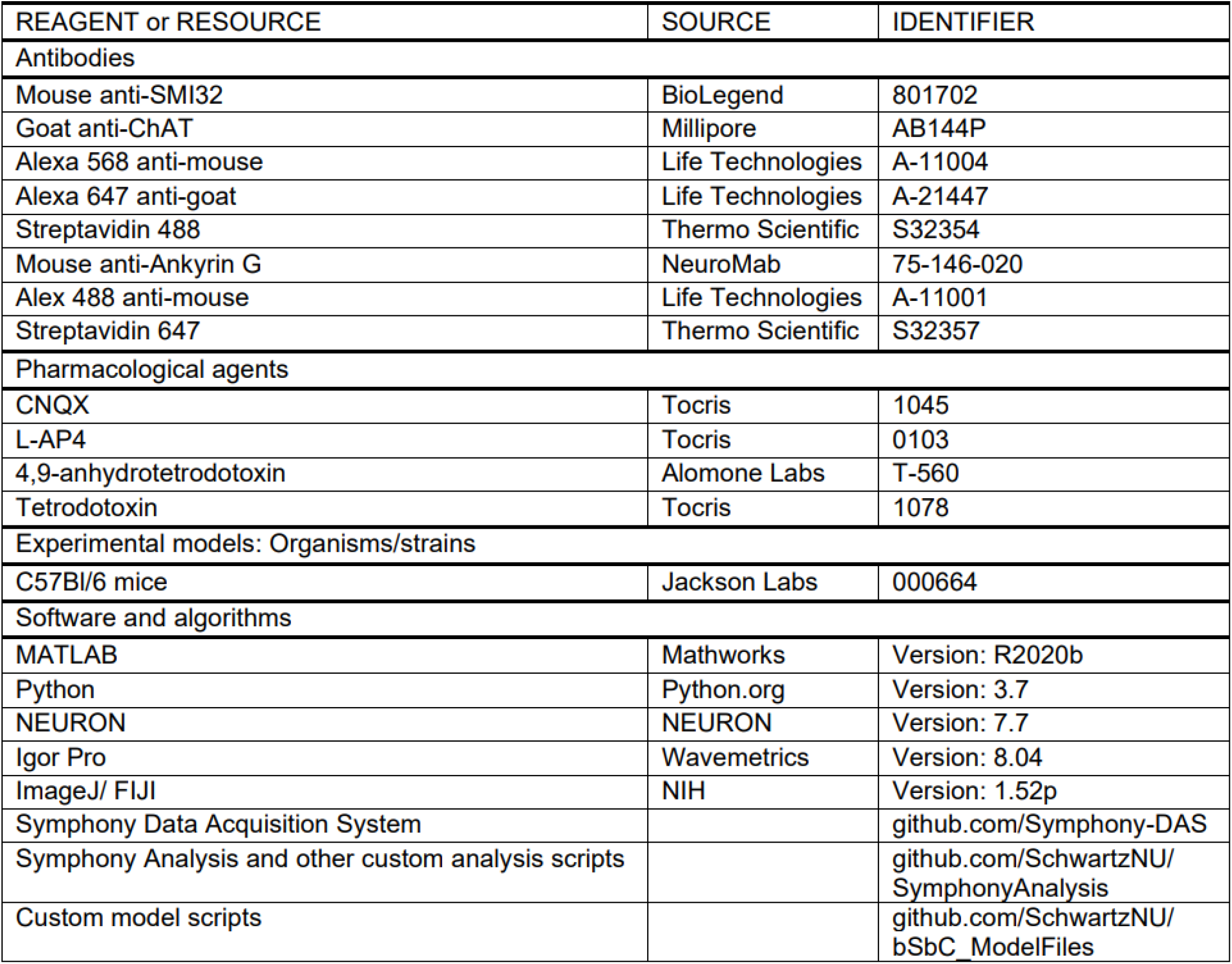

| REAGENT or RESOURCE | SOURCE | IDENTIFIER |
| --- | --- | --- |
| Antibodies |  |  |
| Mouse anti-SMI32 | BioLegend | 801702 |
| Goat anti-ChAT | Millipore | AB144P |
| Alexa 568 anti-mouse | Life Technologies | A-11004 |
| Alexa 647 anti-goat | Life Technologies | A-21447 |
| Streptavidin 488 | Thermo Scientific | S32354 |
| Mouse anti-Ankyrin G | NeuroMab | 75-146-020 |
| Alex 488 anti-mouse | Life Technologies | A-11001 |
| Streptavidin 647 | Thermo Scientific | S32357 |
| Pharmacological agents |  |  |
| CNQX | Tocris | 1045 |
| L-AP4 | Tocris | 0103 |
| 4,9-anhydrotetrodotoxin | Alomone Labs | T-560 |
| Tetrodotoxin | Tocris | 1078 |
| Experimental models: Organisms/strains |  |  |
| C57Bl/6 mice | Jackson Labs | 000664 |
| Software and algorithms |  |  |
| MATLAB | Mathworks | Version: R2020b |
| Python | Python.org | Version: 3.7 |
| NEURON | NEURON | Version: 7.7 |
| Igor Pro | Wavemetrics | Version: 8.04 |
| ImageJ/ FIJI | NIH | Version: 1.52p |
| Symphony Data Acquisition System |  | github.com/Symphony-DAS |
| Symphony Analysis and other custom analysis scripts |  | github.com/SchwartzNU/<br>SymphonyAnalysis |
| Custom model scripts |  | github.com/SchwartzNU/<br>bSbC_ModelFiles |

## Supplemental Figures

**Figure S1.**
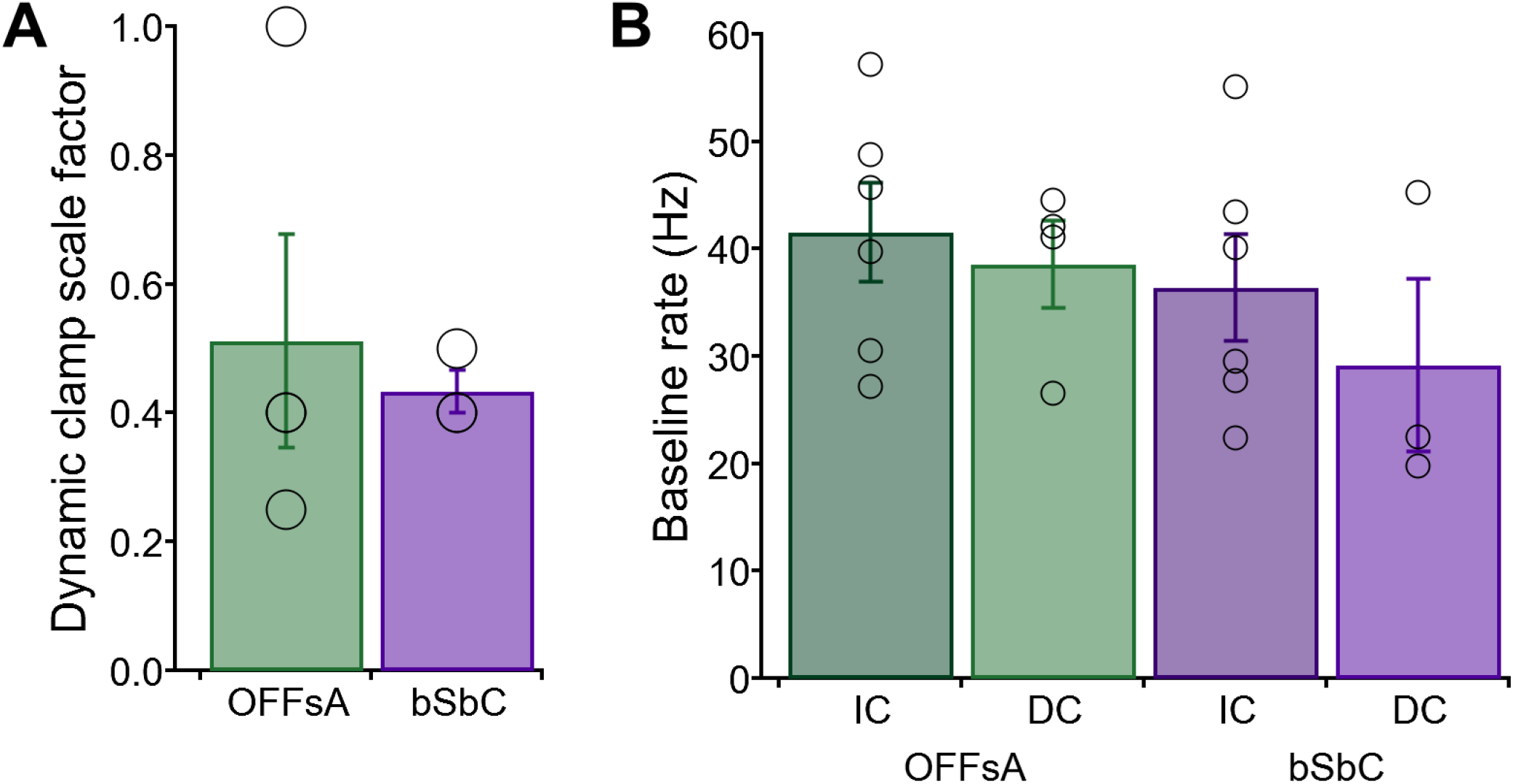
Dynamic clamp parameters are not significantly different between RGC types. **(A)** Dynamic clamp conductance scale factor for recordings in each RGC type (T-test, p = 0.71). OFFsA N = 4, bSbC N = 3. **(B)** Comparison of baseline firing rates in the 2 RGC types in current clamp in the absence of synaptic blockers (IC) and in dynamic clamp with synaptic blockers and baseline conductances applied (DC). OFFsAs, IC N = 6, p = 0.67; bSbCs, IC N = 6, p = 0.45.

**Figure S2.**
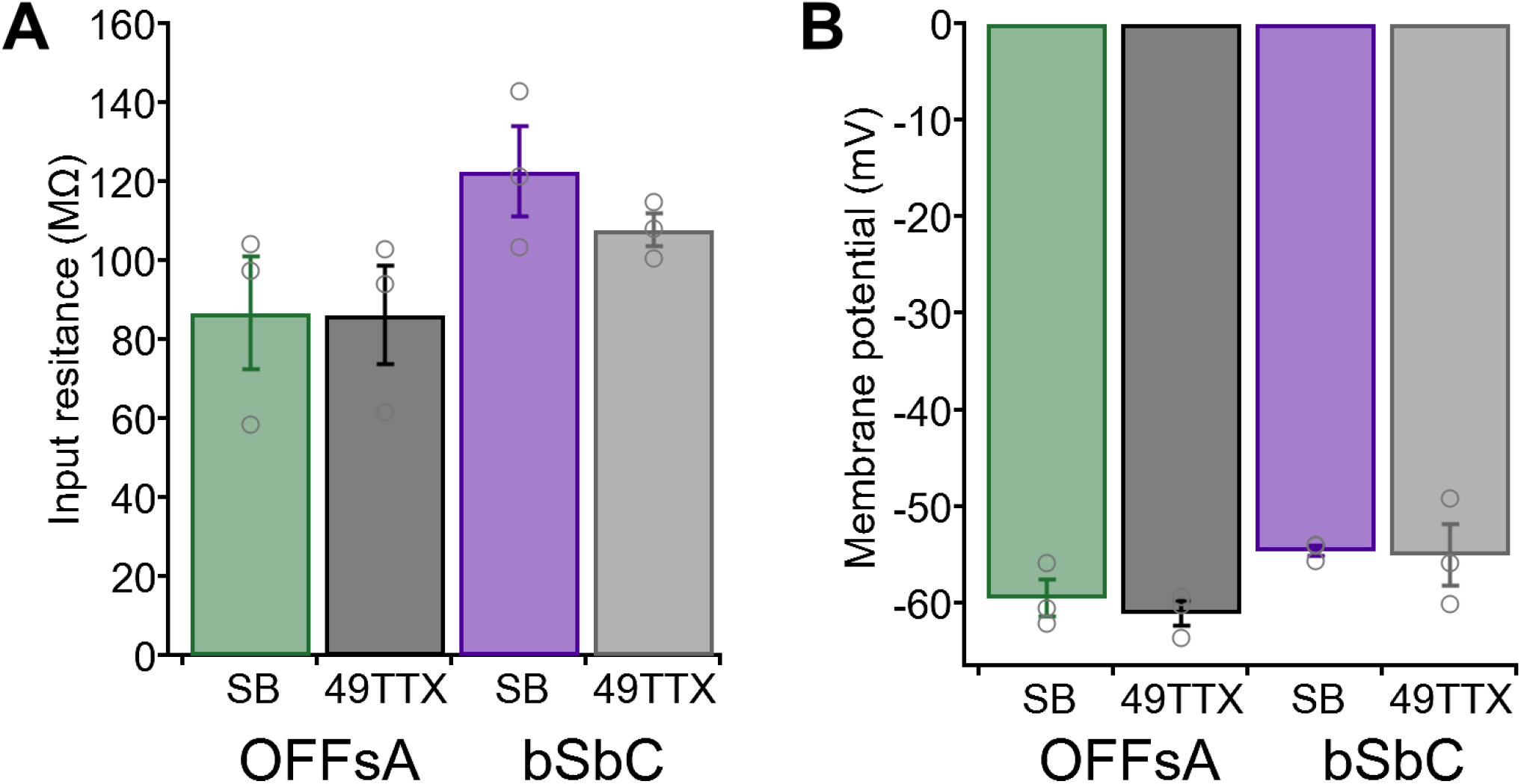
Passive properties are not significantly changed in an Na_v_1.6-specific channel blocker. **(A)** Input resistance for the OFFsA (p = 0.83, paired T-test) and bSbC (p = 0.28, paired T-test) in synaptic blockers (SB) and in SB + 49TTX. OFFsA is *green* for SB and *black* for 49TTX, bSbC is *purple* for SB and *gray* for 49TTX. N = 3 for both cell types. **(B)** Membrane potential for the OFFsA (p = 0.29 paired T-test) and bSbC (p = 0.91, paired T-test) in synaptic blockers (SB) and in 49TTX.

**Figure S3:**
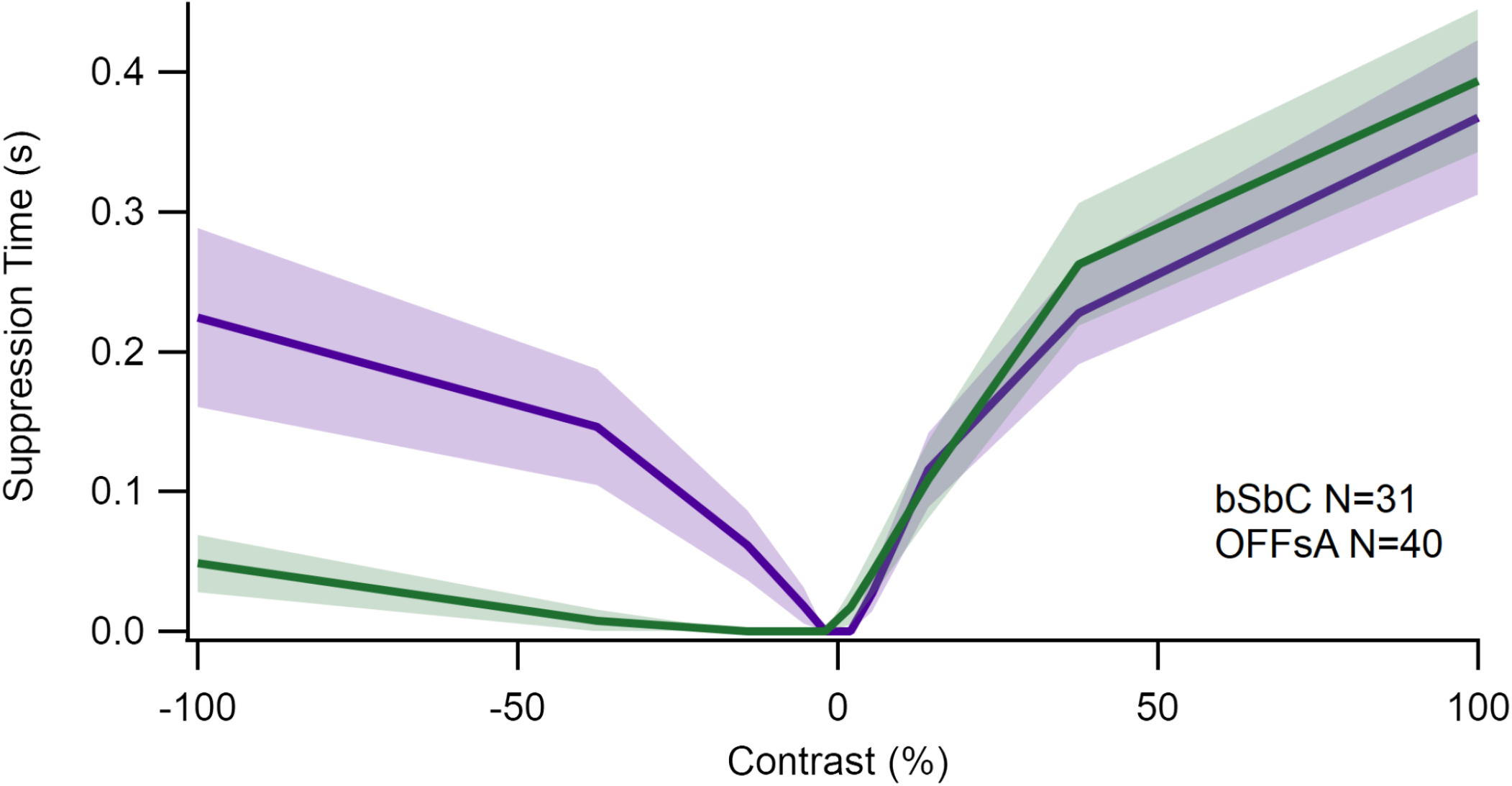
OFFsA and bSbC suppression times differ. **(A)** Suppression time per contrast step for the OFFsA and bSbC. OFFsA N = 40, bSbC N = 31. Suppression time was calculated as the largest inter-spike interval during the stimulus period.

**Table S1:** Compartmental model parameter inputs. For shared values, numbers in parentheses show the computed value for OFFsA RGCs and bSbC RGCs respectively. The value used in the model is the average of the values for the two RGCs.

| Parameter | OFFsA value | bSbC value | Source |
| --- | --- | --- | --- |
| Max $G_{Na}$ (nS) | 270 | 170 | Computed from maximum measured action potential slope, capacitance, and $Na^+$ driving force. |
| AIS Length ($\mu m$) | 20.5 | 14.5 | AnkG staining (**Fig. 8**). |
| Nav1.6 Ratio | 0.4 | 0 | Estimated from reduction of maximum action potential slope in Nav1.6 blockage (**Fig. 7**). |
| Shared values |  |  |  |
| Capacitance (pF) | 58<br>(OFFsA 65, bSbC 52) |  | Computed from the input resistance of the cell times the tau from an exponential fit to the membrane voltage in response to hyperpolarization. |
| Dendritic Length ($\mu m$) | 513 (673, 352) | | Computed from measured input resistance and capacitance after subtracting the membrane area of the other compartments. |
| Soma Diameter ($\mu m$) | 17.5 (18.5, 16.5) | | Light microscopy (**Fig. 1**). |
| Hillock Length ($\mu m$) | 20.75 (23, 18.5) | | AnkG staining (**Fig. 8**). |
| Axon Length ($\mu m$) | 1000 | | Arbitrary (axon is not a propagating compartment in the model). |
| Max $G_K$ (nS) | 360 (440, 280) | | Computed from minimum measured action potential slope (falling phase), capacitance, and $K^+$ driving force. |
| $Na_v$ and $K_v$ channel density ratio: AIS / other compartments | 30 | | Reference (Bender and Trussell 2012). |
| Specific capacitance ($\mu F/cm^2$) | 1 | | Reference (Hille 2001). |
| Axial resistance ($\Omega \cdot cm$) | 200 | | References (Schachter et al. 2010; Abbas et al. 2013). |
| Temperature ($^{\circ}C$) | 32 | | Set in experiments. |
| Membrane resistance ($\Omega \cdot cm^2$) | 3900 | | Computed based on total membrane area and measured input resistance (Freed et al. 1992). |
| $Na^+$ reversal potential (mV) | 30 | | Adjusted down so the spikes were the correct height. |
| $K^+$ reversal potential (mV) | -90 | | References (Hille 2001; Hodgkin and Huxley 1952). |
| Leak reversal potential (mV) | -50 |  | Computed based on measured resting potential. |

**Table S2:**
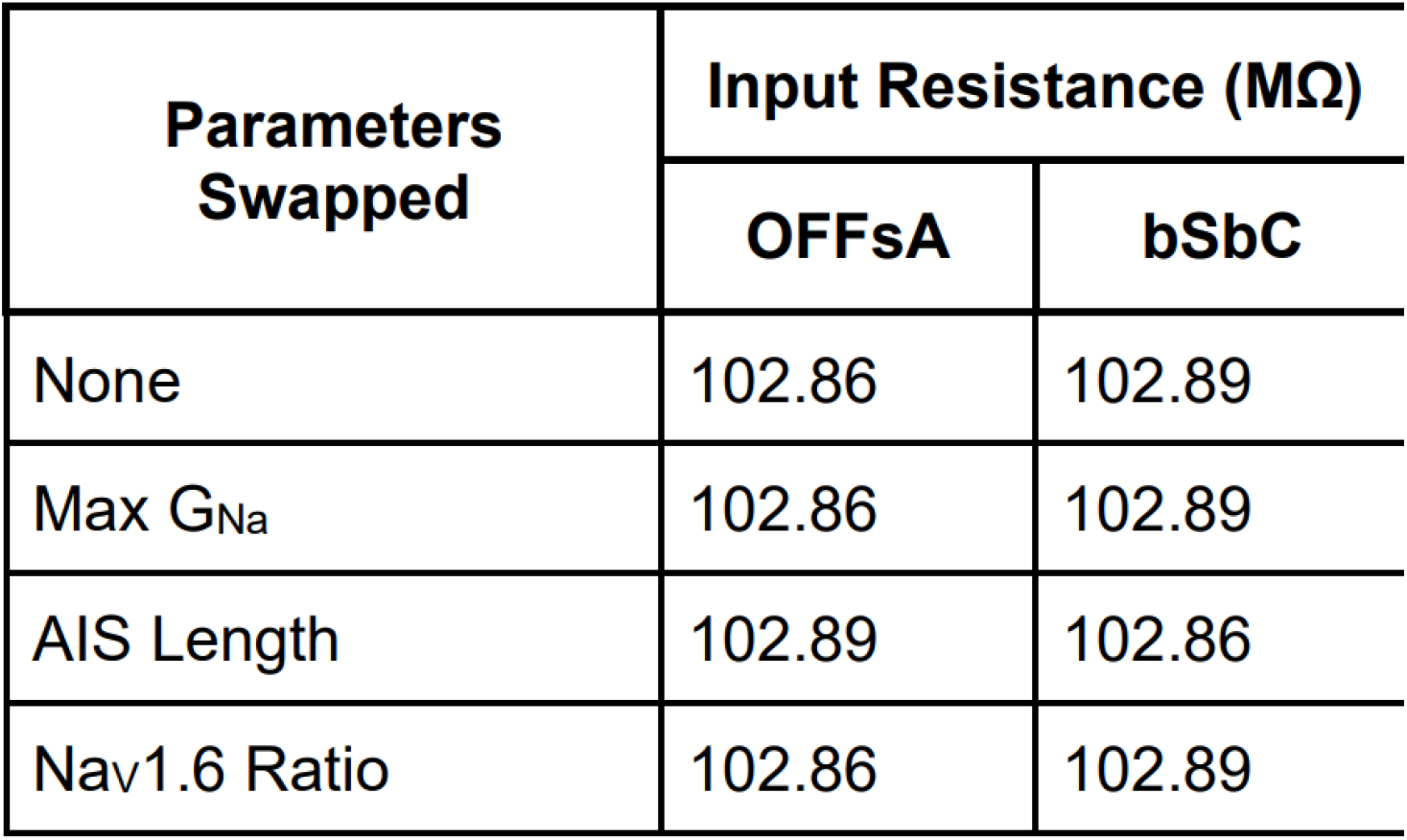
Input resistance for the different models. None indicates that the model has only OFFsA or bSbC parameters **Figure 9C**, while the swapped parameter values correspond to the models in **Figure 9D-F**.

